# The complex genetic architecture of shoot growth natural variation in *Arabidopsis thaliana*

**DOI:** 10.1101/354738

**Authors:** Marchadier, Hanemian, Tisné, Bach, Bazakos, Gilbault, Haddadi, Virlouvet, Loudet

## Abstract

One of the main outcome of quantitative genetics approaches to natural variation is to reveal the genetic architecture underlying the phenotypic space. Complex genetic architectures are described as including numerous loci (or alleles) with small-effect and/or low-frequency in the populations, interactions with the genetic background, environment or age… Linkage or association mapping strategies will be more or less sensitive to this complexity, so that we still have an unclear picture of its extent. By combining high-throughput phenotyping under two environmental conditions with classical QTL mapping approaches in multiple *Arabidopsis thaliana* segregating populations as well as advanced near isogenic lines construction and survey, we have attempted to push back the limits of our understanding of quantitative phenotypic variation. Integrative traits such as those related to vegetative growth used in this work (highlighting either cumulative growth, growth rate or morphology) all showed complex and dynamic genetic architecture with respect to the segregating population and condition. The more resolutive our mapping approach, the more complexity we uncover, with several instances of QTLs visible in near isogenic lines but not detected with the initial QTL mapping, indicating that our phenotyping resolution was less limiting than the mapping resolution with respect to the underlying genetic architecture. In an ultimate approach to resolve this complexity, we intensified our phenotyping effort to target specifically a 3Mb-region known to segregate for a major quantitative trait gene, using a series of selected lines recombined every 100kb. We discovered that at least 3 other independent QTLs had remained hidden in this region, some with trait- or condition-specific effects, or opposite allelic effects. If we were to extrapolate the figures obtained on this specific region in this particular cross to the genome- and species-scale, we would predict hundreds of causative loci of detectable phenotypic effect controlling these growth-related phenotypes.

## Introduction

Fine-tuning plant growth throughout development and in response to environmental limitations is a decisive process to optimize fitness and population survival in the wild. As a sessile organism, plants have to cope with environmental fluctuations and evolved a wide range of responses. This is well illustrated by their great phenotypic plasticity and their ability to colonize very diverse habitats, through intraspecific genetic diversity as revealed in most pathways [1]. Aerial and below-ground growth represent a balance between resource investment in the structures and resource acquisition (respectively photosynthesis and water / nutrient uptake). Thus, growth is a highly complex trait controlled by many genes with constitutive or more specific roles depending on developmental stage, tissue, timing, environment… [2–7]. In this context, plant growth can be considered as a model complex trait to increase our knowledge in the genetics of adaptation/evolution, as well as to improve plant performance.

Forward mutant analysis plays a central role in plant biology to blindly identify gene functions associated with a phenotype [8], but sometimes remains limiting to reveal genes with modest phenotypic effect, or when addressing genes from redundant families. With regard to growth and stress tolerance, these limitations are likely to be relevant given the multigenic nature of growth phenotypes, the low mean effect at each locus and/or epistatic interaction they involve [9, 10]. Thus, the use of naturally-occurring variation through quantitative genetic approaches designed to map quantitative trait loci (QTLs) is interesting notably to complement the search for alleles selected during evolution which may not be brought out with classical loss-of-function approaches. Linkage mapping and genome-wide association lead to the identification of large amount of alleles involved in intraspecific phenotype variation from different plant species [1, 11].

With the drop of sequencing and genotyping costs, phenotyping clearly is the limiting factor for quantitative genetics approaches [12]. However the complexity of the genetic architecture of a given trait, which depends on the contribution and the number of loci controlling a trait and their interactions with the genetic background and the environment, has direct consequences on how much phenotyping remains limiting. Highly heritable traits with a limited number of contributing loci (in a given segregating material, or at the species scale) are more likely to be well understood than more complex traits. For instance, a large part of the phenotypic variation for flowering time in *Arabidopsis thaliana* maps to a limited number of loci [13–16], including *FRIGIDA* and *FLC* genes [17–19] and thus has a relatively simple genetic architecture, although many more loci make smaller contributions -at least in some environments- and allelic heterogeneity also interferes [15, 16, 20, 21]. By contrast, traits like fitness or growth can be expected to have a more complex basis as they integrate many upstream traits, and consequently many genes, each prone to residual variation and heterogeneity. Smaller contributions from individual loci means that, although one can still estimate total heritabilities, the accuracy and throughput of phenotyping will be limiting to confirm individual QTLs’ contributions. Heritabilities for flowering time-related traits will often be above 80%, while biomass accumulation or fitness’ heritabilities are essentially found in the range 20-60% [22–27].

Another factor that will influence the genetic complexity of a trait is its response to the environment through phenotypic plasticity [28]. Part of the environmental fluctuations may be controlled in an experimental design, while another part may contribute to the residuals / noise. Whether the sensitivity of a pathway or trait to the environment depends on the number and architecture of the contributing loci remains an open question, however the relationships between higher plasticity and lower heritability are described [26]. Water availability is an environmental factor that varies through space and time and shows great heterogeneity which certainly constrains plant growth and shapes plant distribution in nature and in agricultural systems. Prevalence of drought episode is expected to increase with global climate change making the understanding of plant response to drought one of the major challenge of the next decades [29, 30]; this includes deciphering the genetic basis for variation in mechanisms such as drought escape, avoidance and tolerance [31]. Hence, this environmental parameter is definitely a good candidate to understand the genetic architecture of GxE. However, drought is both difficult to control and hard to predict, because of interactions with almost all other factors in the environment (temperature, air flow, light…) and interplay with other constraints (especially nutrient-related or osmotic). The development of robotic phenotyping tools throughout the community makes it now feasible to acquire traits on hundreds or thousands of plants in precisely controlled and reproducible conditions [32–35], pushing a bit further one of the main limitation for a better decomposition of the genetic architecture of these complex traits.

Still, regarding plant growth variation in nature, mainly genomic regions with relatively large effect were identified in Arabidopsis and were often related to development, immunity or major hormones (for instance [36–43]). A limited number of non-theoretical studies seem to confirm that many genes with smaller effect –potentially involving epistasis and linked loci– would be responsible for part of the phenotypic variation of such complex traits (for instance [44–46]).

Here, we undertook a precise analysis of plant growth genetic architecture under both optimal watering condition and mild drought stress, using a classical linkage mapping approach on 4 biparental segregating populations [47]. As a dynamic trait, we chose to follow growth during the vegetative phase using a high-throughput phenotyping robot (the *Phenoscope* [34]; https://phenoscope.versailles.inra.fr/) to map major- to small- effect QTLs as well as their interaction with drought stress. Zooming in on the loci, we use near-isogenic lines to validate these QTLs and reveal in more detail the genetics behind a single QTL peak. We then focus more precisely on a region where a major Quantitative Trait Gene (QTG) is segregating (= *CRY2*, a known polymorphic actor with major pleiotropic phenotypic consequences), and show that other loci with additive or opposite effect are also present in its vicinity, illustrating the complexity of growth genetic architecture.

## Results

### Characterization of shoot growth-related phenotypes

The four RIL sets used in this study (BurxCol, CvixCol, BlaxCol, YoxCol) were conducted under well-watered (WW) and moderate water deficit (WD) conditions on our high-throughput phenotyping platform. Ensuring that growth occurs in a highly controlled and homogeneous environment, the Phenoscope records a number of image-based quantitative traits describing shoot development (@Figure 1). Taking daily pictures gave access to cumulative (Projected Rosette Area, PRA) and dynamics (Relative Expansion Rate, RER) growth parameters for individual plants (@Figure 1C&D) as well as other descriptive or derived traits (rosette morphology and RGB colour components) [48]. A principal component analysis (PCA) was performed using all picture-based phenotypes at the final day of the experiment, 29 Days After Sowing (DAS; hence ‘PRA29’ etc) and relative expansion rate calculated between 16 and 29 DAS (RER16-29; @Figure 2). The first axis explained a major part of the total variance, essentially through final rosette size (PRA29) and expansion rate (RER16-29). However, PRA29 and RER16-29 variables were not perfectly correlated, with genotypes exhibiting moderate PRA29 despite high RER16-29. The red (Red29) and green (Green29) components colour phenotypes mainly contributed to the second axis and were positively correlated, and both were negatively correlated with compactness at 29DAS (Compactness29) which was the main trait contributing to the third axis. Individual projection showed that the first axis strongly structured the individuals according to the watering treatment (WW versus WD) while the axes 2 and 3 represented cross (RIL set) effects, differentiating CvixCol and BurxCol (axis 2) and BlaxCol (axis 3) from the other RIL sets. PRA29, RER16-29 and Compactness29 are complementary growth phenotypes that were investigated further in this study to quantify different aspects of shoot development variation: final projected rosette area is a cumulative proxy for biomass and photosynthetically-active surface, rosette compactness is an informative parameter -rather stable through time- describing the rosette morphology, and relative expansion rate highlights the dynamics of growth.

**Figure 1:**
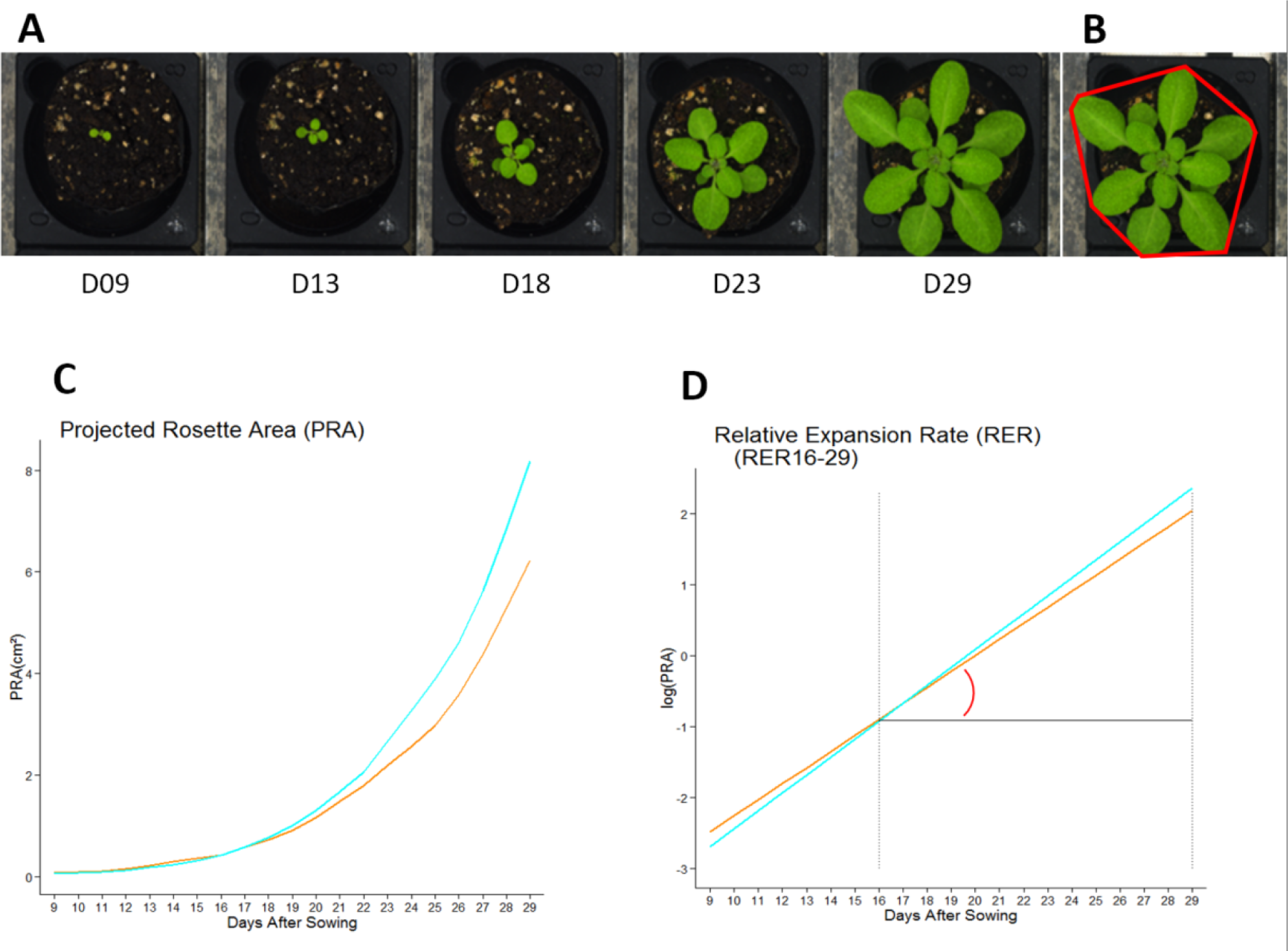
Extracting phenotypic information from zenithal images. A: Pictures of plants on the Phenoscope at different selected days after sowing B: Representation of the rosette-encompassing convex hull (red shape) used to calculate the compactness (here illustrated at day 29) C: Growth kinetics obtained by extracting the projected rosette area from the daily images. D: Relative Expansion Rate calculated between two dates (for instance RER16-29 was integrated from day 16 to 29) C and D: Blue: ‘WW’ = Well Watered (soil water content at 60%); Orange: ‘WD’ = Water Deficit (soil water content at 30%)

**Figure 2:**
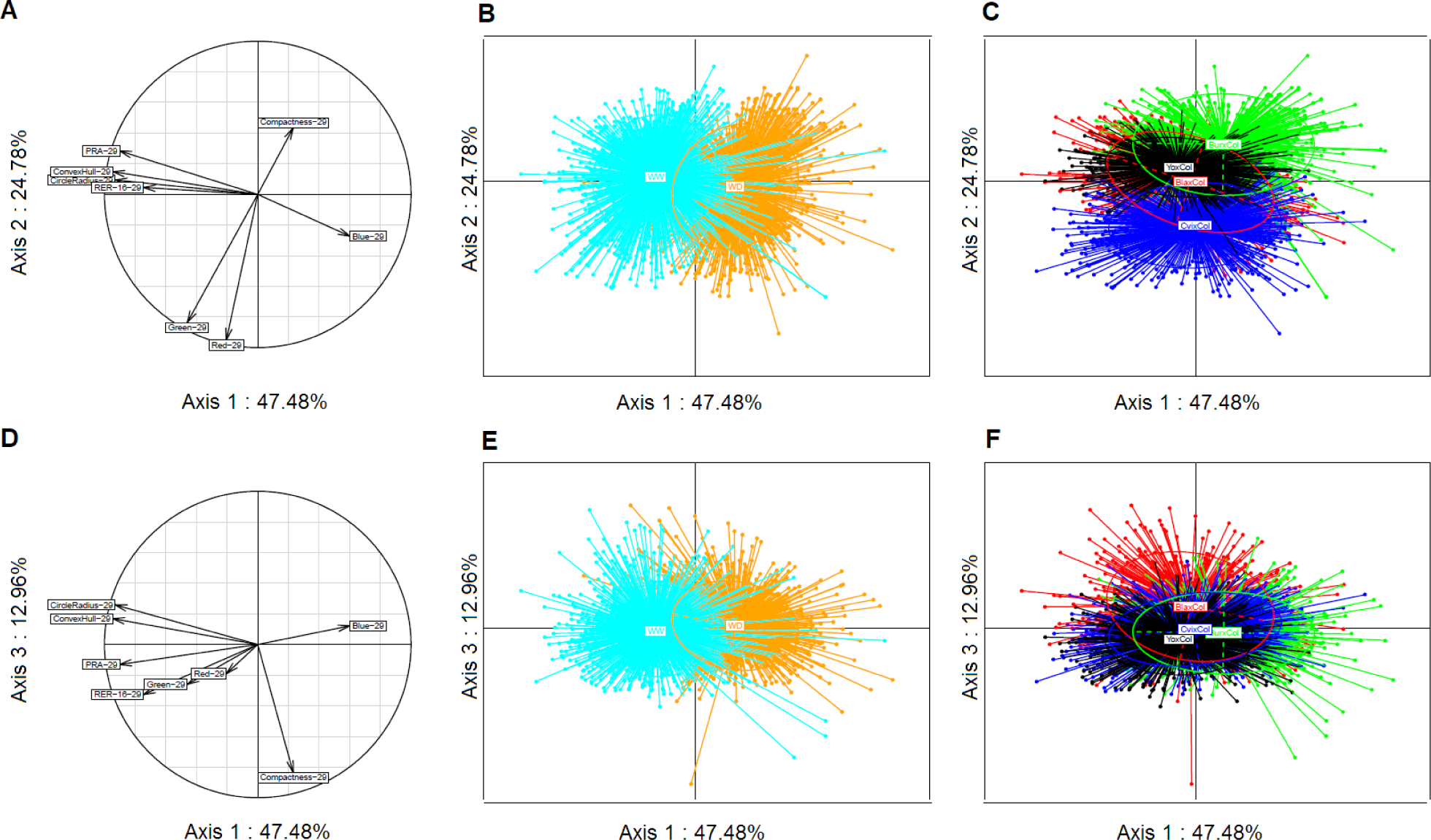
Principal component analysis of phenotypic traits. A to C show the results of the PCA based on plan 1-2. D to F show the results of the PCA based on plan 1-3. A and D: circles of correlations with projected variables on the 2D plan. B and E: individual lines (RILs) projected on the 2D plans, clustered and colored according to the watering conditions as indicated. C and F: individual lines (RILs) projected on the 2D plans, clustered and colored according to the RIL set as indicated.

Phenotypic distribution among the RILs compared to their parents (@Suppl. Figure S1) revealed extensive transgressive segregation for most of the traits and the crosses studied. As expected, mild drought stress (WD) condition impacted the distribution of the RILs for PRA29 and RER16-29 with generally reduced values for both traits. Interestingly, compactness distribution was much more robust to stress, which indicated that overall the morphology of the rosette is less affected by mild drought. In order to estimate the part of phenotypic variation that is explained by genetic factors for each traits, heritabilities were calculated (@Suppl. Table S1) and were essentially low for RER (except in BurxCol where they were high), intermediate for cumulative PRA and generally high for Compactness (this trait is certainly less sensitive to shifts in developmental stage that could be induced, for instance, by small changes in germination time). Overall, heritabilities were also lower in stress conditions than in control, as if the stress was inducing noisier phenotypic variations.

According to ANOVA analyses (@Suppl. Table S2), all traits and RIL sets showed significant variation according to both the watering condition (WD versus WW) and the genotype (within each RIL set), except for Compactness29 response to stress in BlaxCol. There were also weaker but significant biological replicates’ effects (i.e. independent Phenoscope experiments), but less genotype x experiment interactions (with the exception of PRA and RER traits in BlaxCol for instance). PRA is more prone to genotype x experiment interactions than other traits, especially in CvixCol and YoxCol. Genotype x condition interactions are often milder than genotype or condition effects, and overall compactness –or YoxCol– show much less genotype x condition interactions than other traits/sets.

The phenotypic values of each RIL were then corrected for inter-experiment differences (indicated by the significant biological replicates’ effect).

### QTL mapping-based shoot growth genetic architecture

Our experimental design allowed the identification of many QTLs for all combinations of traits, conditions and RIL sets, and also for the GxE interaction term using genotype x condition effects from the ANOVA model for each trait (@Figure 3 and @Suppl. Table S3). Globally 112 QTLs were identified all along the genome when conditions are studied independently (62 under WW + 50 under WD) likely corresponding to at least 18 independent loci, yielding an average of ˜4.7 QTLs per modality of cross x trait x condition. QTL hotspots across RIL sets and traits were identified for instance at the beginning of chromosome 1, at the bottom of chromosome 2 and 5. These hotspots include very highly significant QTLs with LOD scores above 10, and up to 32. Chromosome 3 appeared to show less significant QTLs in all crosses, especially for PRA29 and RER16-29. Individual QTL contributions to phenotypic variance (R²) ranged from 1 up to 30%, with the typical L-shaped distribution of effect. Using empirical significance boundaries according to the observed distribution of QTLs effects, ˜10% of the QTLs could be considered as showing major effects and significance (R2>10% and/or LOD>15); ˜25% of the QTLs could be considered as showing intermediate effects and significance (5%<R2<10% and/or 7<LOD<15); the remaining 2/3rd of the QTLs could be considered as showing minor effects and significance R2<5% and/or LOD<7). Many more potential QTLs not listed here were only suggestive with LOD score peaks just below our threshold (<2.4 LOD).

**Figure 3:**
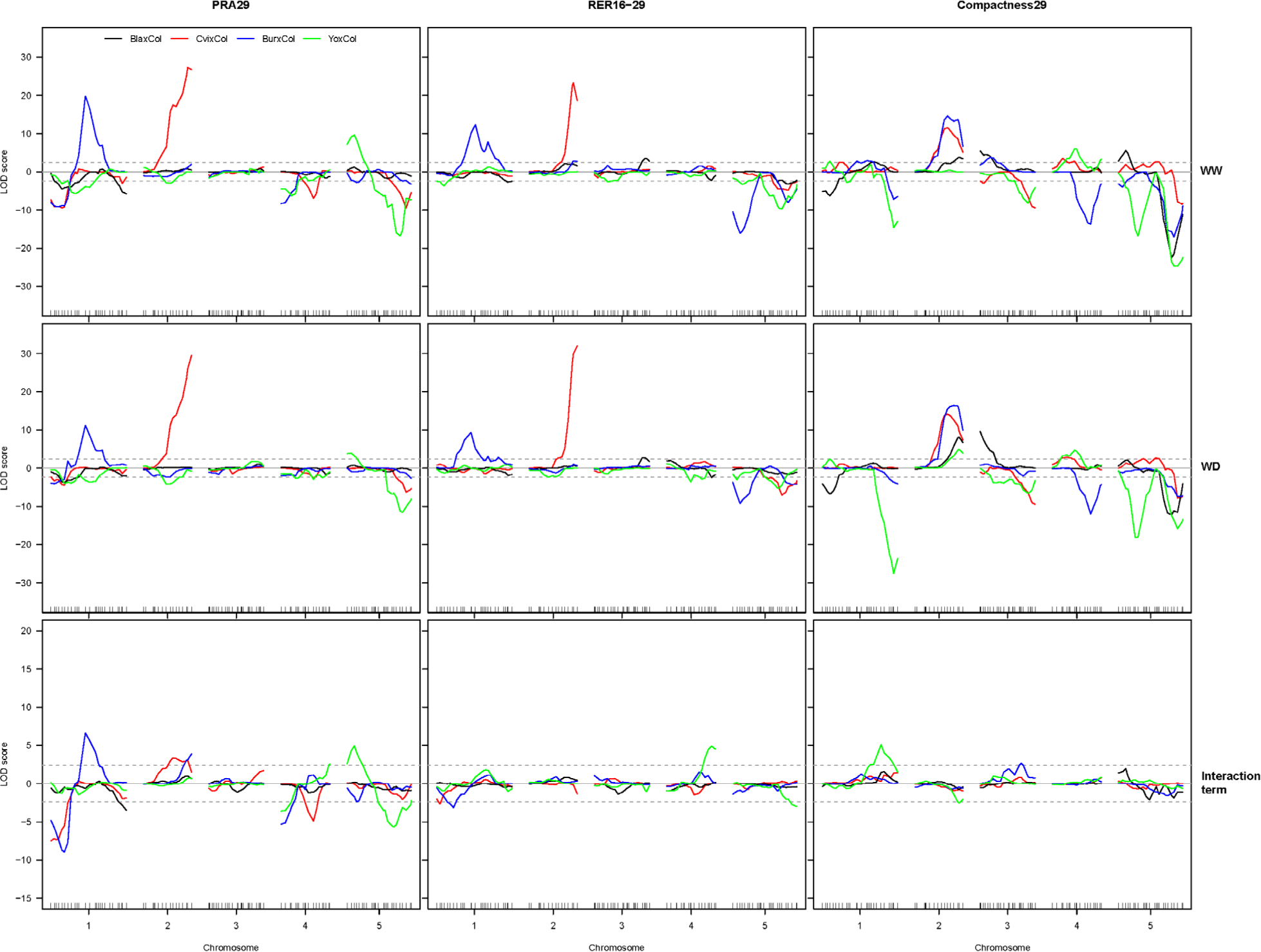
QTL mapping for 4 RIL sets, 3 traits and 2 conditions. QTL maps obtained from four RIL sets (BlaxCol in black; CvixCol in red; BurxCol in blue; YoxCol in green) using three phenotypes (PRA29, RER16-29 and Compactness29), generated independently in two conditions (top panels: WW; middle panels: WD) and from the genotype x environment interaction term in the model (bottom panels: Interaction term). LOD score values above (below) the X axis indicate that the Col allele increases (decreases) trait value with respect to the other parental allele. Significance threshold (LOD=2.4) is shown as dotted lines. The parameters of the significant QTLs detected are listed in @Suppl. Table S3.

Most of the QTL profiles are stable across the 2 watering conditions, especially for the major-effect loci. However, QTLs specific for one condition were detected, e.g for RER16-29 under WD in YoxCol on chromosome 4, and at the top of chromosome 1 under WW. We also mapped QTLs for the interaction term with the drought treatment, yielding another 19 QTLs (@Figure 3 and @Suppl. Table S3). These essentially emphasize large effect QTLs showing a modulation of their effect in response to stress (especially for PRA, see chromosome 1 and 5), with no real new loci revealed. For RER16-29 in YoxCol, this would confirm the interaction of the above-mentioned locus on chromosome 4 (although its exact location is questionable), but not for the one at the top of chromosome 1. There may be some power issues when comparing across conditions due to lower heritabilities of traits under WD.

Although derived from PRA29, Compactness29 showed an independent genetic basis, as exemplified by the major peaks on the bottom of chromosome 4 in BurxCol or 5 in BlaxCol. Although contributing to RER16-29, PRA29 does not always share the same contributing loci, for instance on the top of chromosome 1 in CvixCol and BurxCol. Other more complex cross x trait patterns are apparent, like at the bottom of chromosome 2 where a major QTL for Compactness29 in three of the four RIL sets seems to colocalize with a significant PRA29 and RER16-29 QTL (in the same direction), but only in one cross. It may be that these Compactness29 QTLs are actually independant in each RIL set.

A two-dimensions search for epistatic interactions was performed across all traits, conditions and RIL sets (@Suppl. Figure S2). Overall, the BurxCol and CvixCol RIL sets showed more significant epistasis compared to the two other sets. Interestingly, pairwise interactions controlling growth phenotypes are overall quite different depending on RIL set and growth phenotypes. Shared epistasis effects are potentially detected in both watering conditions: they appear as symetrical across the diagonal on @Suppl. Figure S2. One of the most significant epistatic interaction was observed between 2 loci on the top and bottom of chromosome 4 in the BurxCol cross (@Suppl. Figure S2; highest significance for RER16-29 in WW). Positions and directions of effect match perfectly with the previously published SG3 x SG3i interaction known to segregate in this cross [49]. Based on its significance, another relevant interaction (*p*-value=0.01 in WW and *p*-value=0.03 in WD) was observed for PRA29 in the BlaxCol cross between the bottom of chromosomes 4 and 5 (@Figure 4). The effect of the bottom of chromosome 5 QTL on PRA29 is observed only when RILs carry Bla alleles at the bottom of chromosome 4. As a consequence of this epistasis, these QTLs appear barely significant in single QTL scans (@Figure 3).

**Figure 4:**
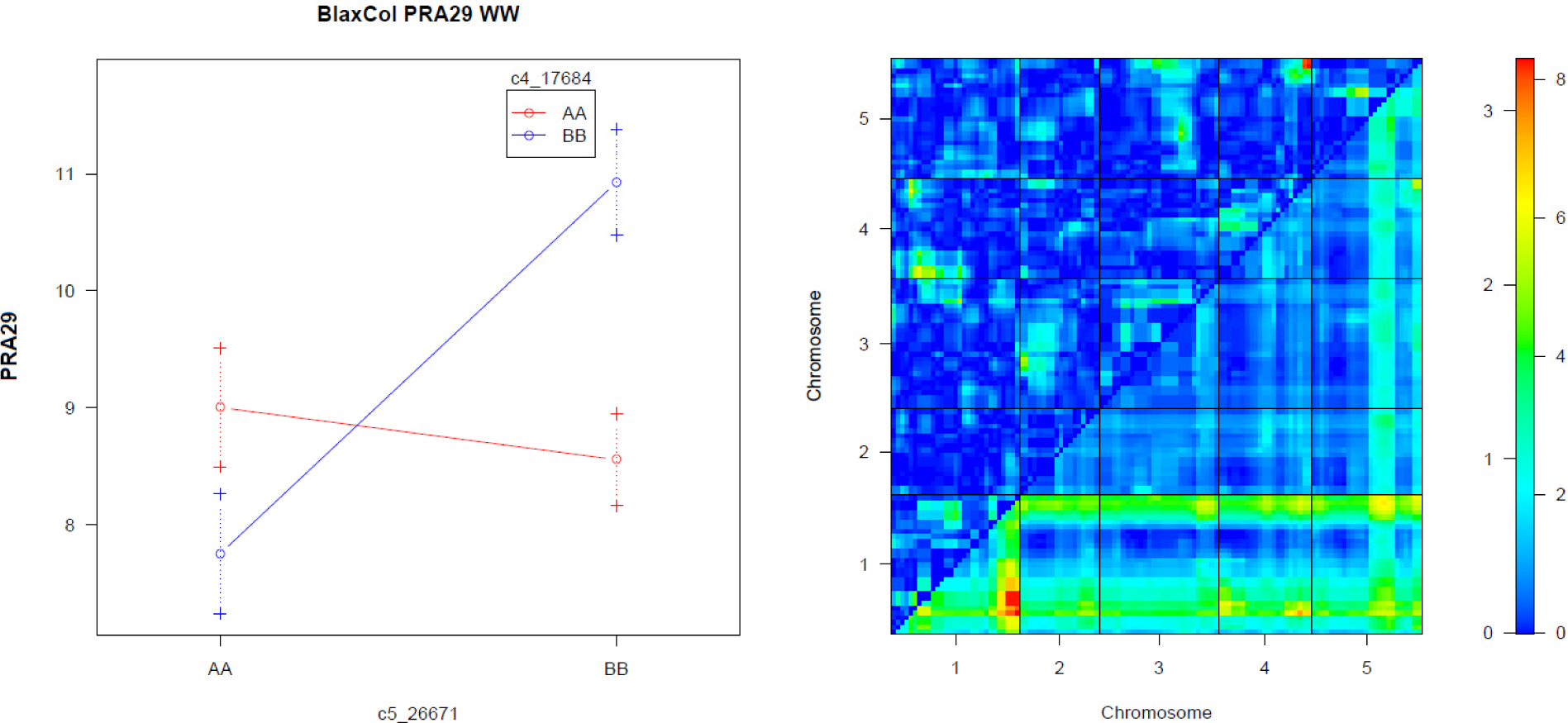
A significant epistatic interaction for PRA29 in BlaxCol. A significant epistatic effect on PRA29 between the bottom of chromosomes 4 and 5 segregating in the BlaxCol RIL set. Results shown in WW condition. The interaction plot (left panel) represents the average RIL phenotype (in cm2) for each allelic combination (‘AA’ = homozygous Col, ‘BB’ = homozygous Bla). Right panel: heatmaps representing the LOD score of the full model with additive and interaction effects (triangle below diagonal) and that of the interaction effects only (triangle above diagonal) for PRA29 in BlaxCol. The color scales for significance (LOD) are indicated on the right.

We also performed dynamic QTL detection on daily-recorded traits (PRA and Compactness) to reveal the evolution of QTL effect throughout the experiment on the Phenoscope: these interactive QTL profiles can be accessed at http://www.inra.fr/vast/PhenoDynamic.htm Most of the PRA QTLs observed after 29 days of growth correspond to locus that become gradually significant across the experiment and are essentially not time-specific. These are most likely contrasted allelic effect on growth that cumulate their effect over time. There are only a few exceptions for PRA, like the bottom of chromosome 5 locus segregating in BlaxCol which remains significant only until 13DAS and thereafter is canceled out. For Compactness, the picture is rather different with numerous examples of QTLs that are essentially significant around specific time-points, even sometimes in successive waves of significance (BlaxCol, bottom of chromosome 5: QTL peaking at Days 11, 17 and 27 –providing that this is a unique locus).

In order to take advantage of the different crosses to Col-0 in our experimental setup, QTL mapping was performed on the whole dataset using MCQTL tool to compare allelic effects in a multicross design and potentially reveal shared QTLs. The combined QTL maps obtained (@Figure 5 for PRA29 and @Suppl. Figure S3 for RER16-29 and Compactness29) highlight 10 independent loci, including at least 4 regions with contrasting allelic effect on PRA29: for instance, the middle of chromosome 1 region shows contrasted phenotypic consequences in different crosses, particularly when comparing Bur and Yo alleles (with respect to Col). At this scale, it can be difficult to distinguish between different alleles at the same QTL and different QTLs. Conversely, combining the information of multiple crosses sometimes allows to predict narrower QTL intervals than with the initial QTL mapping, enabling the detection of distinct linked loci, for instance for PRA on the bottom half of chromosome 2; a location where the dynamic QTL analysis for PRA in CvixCol was already showing signs of 2 different segregating loci with slightly distinct dynamics over time. Another striking example is for Compactness on chromosome 5, with neighbouring QTLs predicted to show opposite allelic effects (e.g. BlaxCol).

**Figure 5:**
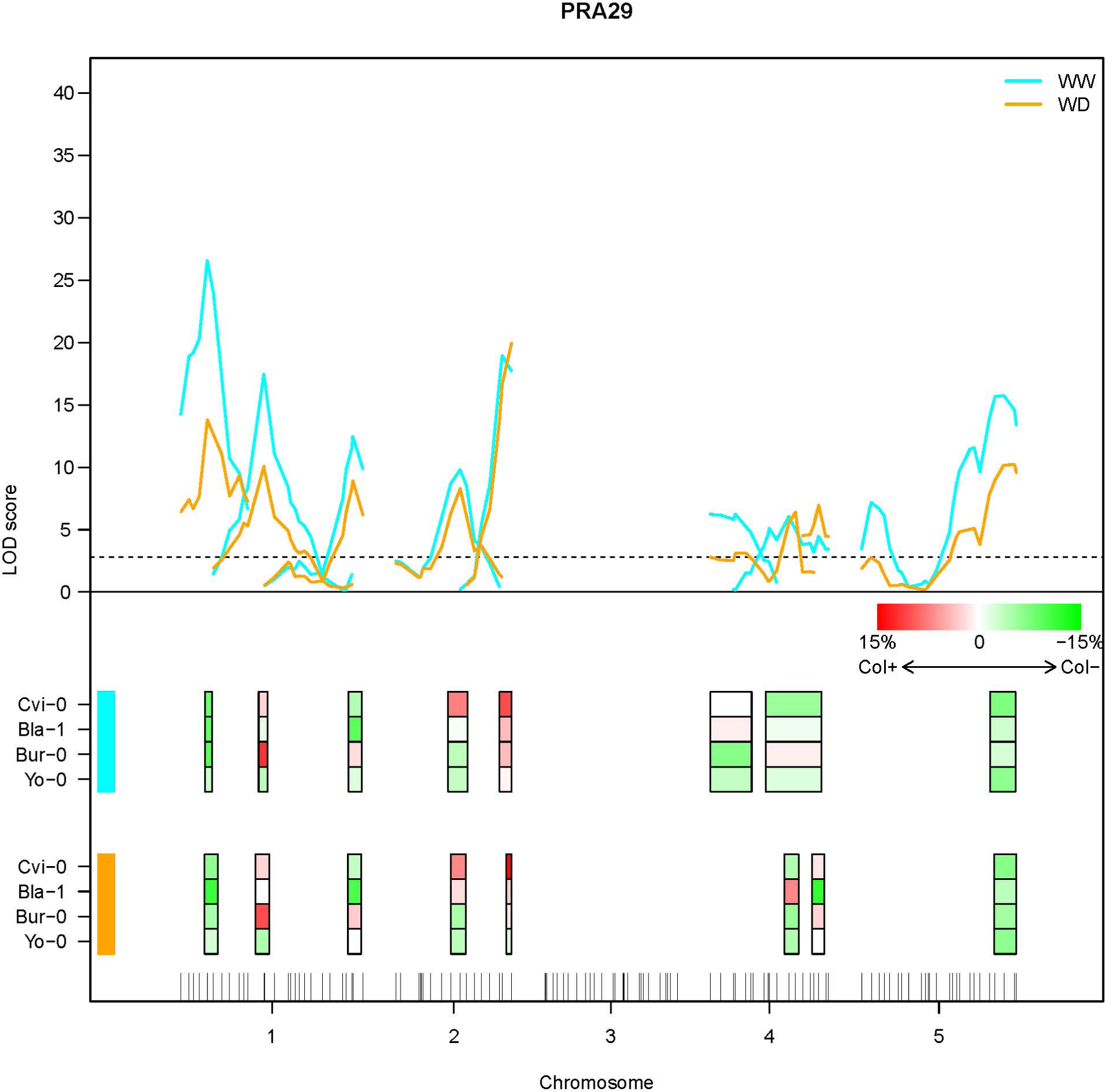
Multi-cross QTL analysis for PRA29. The upper panel of the figure represents the LOD scores calculated by the MCQTL method along the genetic map in WW (blue) and WD (orange) conditions. Each QTL is depicted by a LOD score profile which peak is represented by colored boxes on the lower panel of the figure (WW conditions above and WD conditions below). The color of the box indicates the deviation of the trait value for the Cvi, Bla, Bur or Yo alleles relative to the Col allele as indicated. Chromosome 3 shows no significant combined QTLs for PRA29. This figure shows results obtained for trait PRA29 only; other traits are in @Suppl. Figure S3.

### Near isogenic lines-based shoot growth genetic architecture

To confirm and investigate further the complexity of the genetic architecture of these traits, we used near-isogenic lines to mendelize QTLs and assess in more details the role of smaller chromosomal regions. Using 81 independent Heterogeneous Inbred Families (HIFs) scattered across the genome or chosen to decompose candidate regions in specific crosses [50], we tested a total of 79 QTL effects from the 24 modalities of RIL set x trait x condition (@Figure 6 for PRA29; @Suppl. Figure S4 for RER16-29 and Compactness29). Globally, 60% of the HIFs with a segregating region covering a candidate region previously identified showed significant effect with consistent direction, thus validating the QTL. Larger effect QTL were more often validated in HIF, with ˜75% of the tested major or intermediate effect QTLs confirmed, compared to ˜50% of the minor effect QTL. Specifically for PRA29 trait in the two conditions studied (@Figure 6), 28 QTL were assessed with at least one HIF, among which 17 (60%) were confirmed. Some (minor-effect) QTL x condition interactions detected in the RIL set were also significantly confirmed using HIF, such as the top of chromosome 5 locus in BurxCol which has significance (PRA29) only under WW.

**Figure 6:**
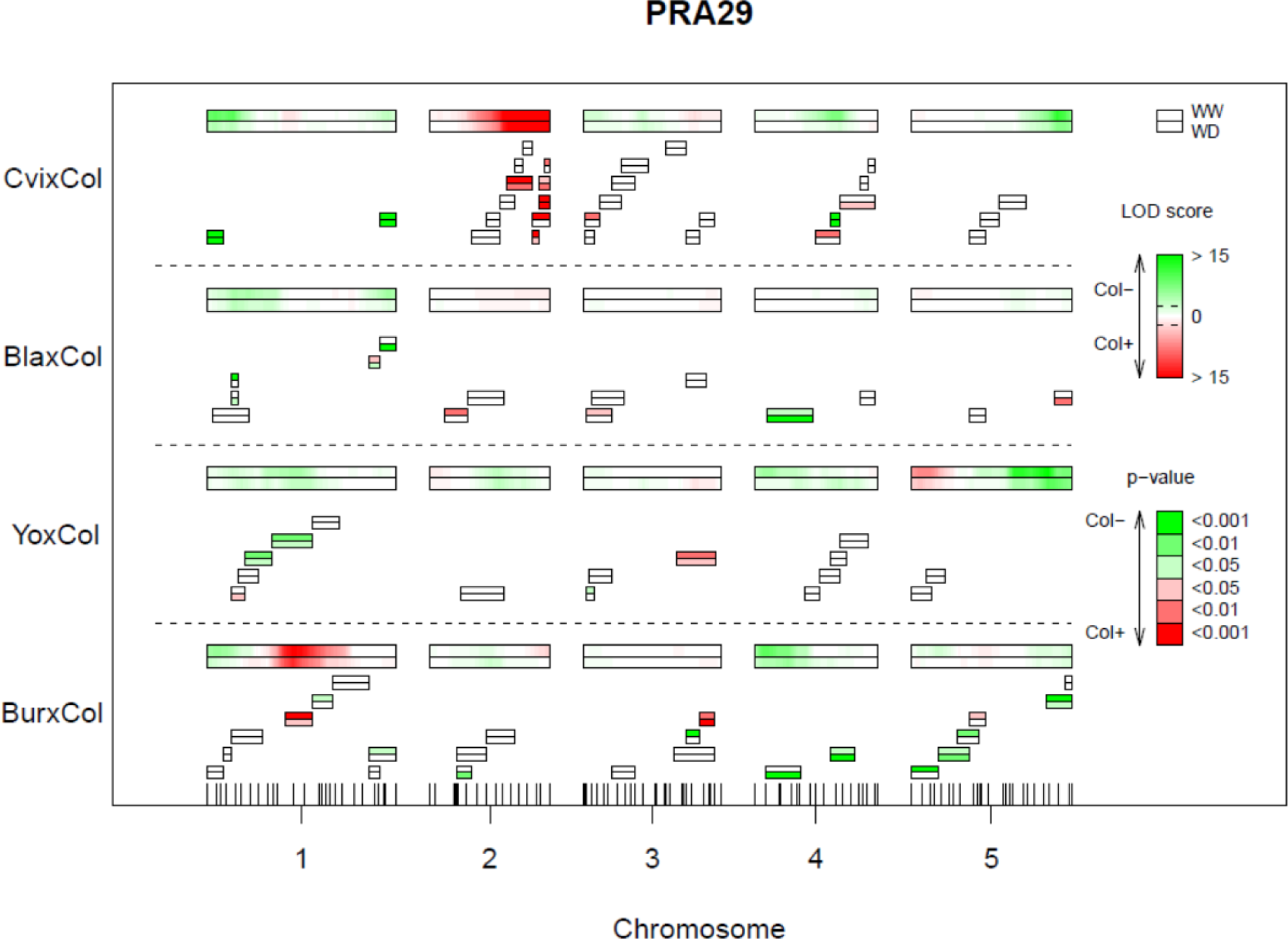
Near isogenic lines-based validation of QTLs for PRA29. QTL mapping results (same LOD score profiles as @Figure 3) are shown as LOD score heatmaps (see color scale on the right) for each RIL set, across long twin boxes representing the 5 chromosomes (the upper frame is for WW, the lower frame is for WD conditions). Below each chromosome are series of short twin boxes representing individual HIFs segregating region, used to test phenotypes associated with specific regions (for each twin box the upper frame is for WW, the lower frame is for WD). The significance and the direction of the difference between Col-fixed HIF lines and alternate parental allele-fixed HIF lines are indicated by the p-value color scale as indicated. Details of each HIF used here are gathered in a deposited dataset [50]. This figure shows results obtained for trait PRA29 only; other traits are in @Suppl. Figure S4.

Many factors can explain that a QTL is not validated in an HIF (beyond an unreliable phenotype): the QTL could be mislocalised by QTL mapping (thus, out of the HIF segregating region), the QTL could be under epistatic interaction with another locus (thus, a specific HIF may not represent the adequate genetic background), or the HIF harbours a more complex genotype than expected at the segregating region, such as a double recombined region (thus, the HIF doesn’t actually allow to test the whole region). This makes the comparison between HIFs difficult, even in the same cross, and negative HIF results should particularly be interpreted carefully. In this context, the rate of QTL validation obtained here is rather satisfying.

We are also interested in positive HIF results that were not predicted by the QTL mapping. This was particularly possible in regions with high HIF coverage and shows that several LOD score peaks were explained by more than one underlying QTL, as also suggested by the MCQTL analysis. A good example lies in the CvixCol cross, where the bottom half of chromosome 4 seemed to control PRA29 as a single locus in the initial QTL analysis (@Figure 3) and is now subdivided in at least 2 independent loci with opposite allelic effects after the HIF analysis. Another example segregates in YoxCol at the top of chromosome 3 where two QTL with opposite-effect on Compactness29 under WW conditions seem to localise closely according to the HIFs (@Suppl. Figure S4) but wasn’t detected at all in the QTL mapping at 29DAS (@Figure 3) and only remained significant at intermediate stages around 16DAS according to the dynamic analysis. The initial QTL mapping may have lacked power (= RILs) to detect these complex patterns, either due to the confusing effect of linked loci, or the confusing effect of epistasis, or both.

Finally, we can occasionally exploit the localisation of the segregating region(s) of the tested HIFs to narrow down the candidate QTL region, like for a PRA29 QTL in CvixCol at the bottom of chromosome 2 which is narrowed down to the extremity of the chromosome and shown to be distinct (= confirmed in non-overlapping HIFs) from another nearby QTL segregating in the same cross (with allelic effect in the same direction), as predicted by the multicross analysis above. Here again the dynamic analysis also helped to distinguish these loci based on their effect through time. Still, the ‘precision’ remains approximately at the Mb level at this stage.

To tackle further the question of the complexity of the genetic architecture at a higher resolution than with simple HIFs, we decided to systematically dissect a region of 3Mb at the very beginning of chromosome 1 in CvixCol: in this region all previous approaches have predicted a single QTL with intermediate to major effect significance on PRA29 (@Figure 3), which was confirmed in an HIF (@Figure 6). This HIF was used to zoom in on the region across 30 bins (stairs) of ˜100kb, defined by successive recombination breakpoints (@Suppl. Table S4) and phenotypically evaluated individually in the same conditions as above (an approach coined ‘microStairs’). We selected this interval to test if a region harbouring a large-effect growth QTL may typically also include independent loci, maybe of smaller effect or hidden by the main QTL. A pairwise comparison of their growth phenotypes allows to test the impact of Cvi versus Col alleles over relatively short physical intervals of expected average size 100kb, with a maximum ˜200kb, depending on recombination breakpoints location and marker interval. We either compared only the 2 successive recombinants (‘stair by stair’), or we took advantage of the support from all pairwise comparisons (‘staircase’) to increase our power to detect QTL (@Figure 7 for PRA29 and @Suppl. Figure S5 for RER16-29 and Compactness29).

**Figure 7:**
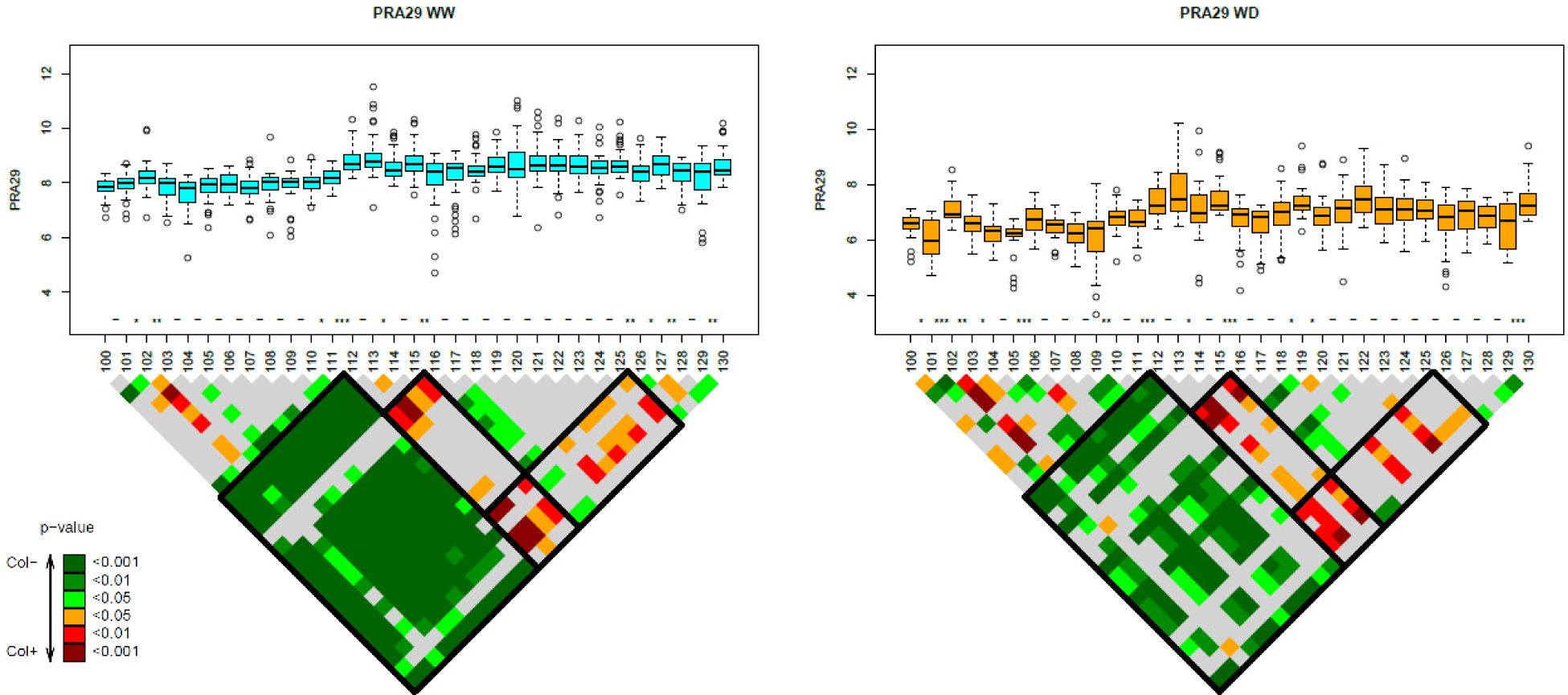
Dissection of a genomic region in CvixCol for PRA29 (‘microStairs’ approach) The upper panels’ are boxplots indicating phenotypic values observed for each of the 31 successive recombinant lines along the 3Mb region studied (details of each line used here are in @Suppl. Table S4). Significant differences between successive lines (stairs) are indicated along the X axis with ‘*’ (p-value < 0.05), ‘**’ (p-value < 0.01), ‘***’ (p-value < 0.001) or ‘-’ for no significant difference. The lower panel shows a triangular matrix gathering the significance of all pairwise comparisons between lines (see the p-value color scale also indicating the direction of the allelic effect). Blocks of significance (‘staircase’) are interpreted and highlighted in black-framed boxes. This figure shows results obtained for trait PRA29 only (in 2 conditions), PRA is expressed in cm2; other traits are in @Suppl. Figure S5.

The strongest phenotypic effect for PRA29 in WW and WD (but also –to a lesser extent– for RER16-29 in WD) was fine-mapped down to the stair between recombinants 111 and 112, covering potentially a physical interval (bin) extending to the maximum between 1.070 and 1.291 Mb (an interval including 54 genes). In the middle of this region lies the obvious candidate gene *CRYPTOCHROME 2* (At1g04400; *CRY2*, a blue-light photoreceptor; @Suppl. Table S5) that is known to harbour a functional variant in Cvi (a single amino-acid change) and impact plant photomorphogenic development especially in short days [51, 52]. This is a good test case of our approach, as it is very likely that *CRY2* is primarily responsible for the growth difference overall observed in this HIF.

Still, on either side of this locus, a few other QTLs also affect growth, underlying a much more complex genetic architecture than expected after the initial HIF results: in both watering conditions, a milder PRA29 QTL, with opposite allelic effect than CRY2, was detected in the stair defined by recombinants 115 and 116 (1.495 to 1.703 Mb positions). Another PRA29 QTL was predicted in the bin 125/126 (2.509 to 2.710 Mb). For Compactness29 specifically under WD treatment, *CRY2* did not seem to be causal and the causative locus for the observed HIF phenotype would most likely be in the bin 118/119 (1.801 to 2.027 Mb); this QTL was not clear from the initial QTL analysis for this trait on Day 29 (@Suppl. Figure S4), but appeared significant earlier in the experiment (cf <u>dynamic analyses</u> for CvixCol around 14DAS and later). There are probably even more loci, especially at the very beginning of the region for PRA29 (stairs between recombinant 101 and 104 at least), but we reach the limits of our experimental design and phenotyping precision to be able to conclude accurately, with either too complex genetic architecture in this region or not enough recombinant lines to robustly support each intervals’ effect.

We then looked for high impact polymorphisms (premature stop codon, frameshift or non-synonymous mutations) likely affecting gene function between Col-0 and Cvi-0 within the most promising bins to find out candidate genes. Because of the numerous non-synonymous changes between these accessions (@Suppl. Table S5), we decided to arbitrarily filter the genes with the criteria of at least 3 non-synonymous mutations to increase our chance to detect genes with altered function or degenerated sequences after loss-of-function mutations. Within the bin 115/116 lie at least 2 interesting candidates (among those with functional annotations):

*URIDINE DIPHOSPHATE GLYCOSYLTRANSFERASE 74E2* (At1g05680; *UGT74E2*) is an auxin glycosyltransferase whose overexpression was shown to modify plant morphology and the size of the petioles, to delay flowering, and to increase drought tolerance [53]. The gene is also known for ample natural variation in expression, including potentially cis-acting variants [54]: http://www.bioinformatics.nl/AraQTL/multiplot/?query=AT1G05680.

*GLUTAMATE RECEPTOR 3.4* (At1g05200; *GLR3.4*) is a calcium-dependent abiotic stimuli-responsive gene [55] expressed throughout the plant and impacting at least lateral root initiation [56]. It harbours several non-synonymous variants in Cvi-0 (compared to Col-0), but also higher expression in Col-0 as shown by several local-eQTLs in different crosses (http://www.bioinformatics.nl/AraQTL/multiplot/?query=AT1G05200) confirmed to be cis-regulated through Allele-Specific Expression assay [57] (reported in @Suppl. Table S6). Within the bin 125/126 lie at least 2 interesting candidates:

AT1G08130 encodes *DNA LIGASE 1* (*LIG1*), a ligase involved in DNA repair, which mutation causes severe growth defects [58]. Its expression is also known to be controlled by a local-eQTL (most likely cis-acting) in LerxCvi: http://www.bioinformatics.nl/AraQTL/multiplot/?query=AT1G08130

AT1G08410 is *DROUGHT INHIBITED GROWTH OF LATERAL ROOTS* 6 (*DIG6*), encoding a large 60S subunit nuclear export GTPase 1 that impacts several developmental processes regulated by auxin, including growth [59].

Further work is required to prove any link between the observed phenotypic variation and these candidate genes.

## Discussion

Owing to its fine regulation throughout development and in interaction with the environment, plant growth represents a highly complex trait potentially controlled by numerous factors and interactions. Little is known on the actual genetic architecture of plant growth natural variation, with essentially a few genes of major effect being identified until now and only a few exceptions of more complex genetics revealed [44]. Association genetics hold great promise to dissect the underlying molecular bases of complex traits [12], however one can wonder if the genetic architecture of highly integrative traits like growth or fitness is amenable to genome-wide association studies (GWAS) at the species scale: GWAS especially lacks power to decompose traits controlled by many loci of small effects, particularly if the underlying alleles have moderate or low frequency in the populations. For instance when exploiting worldwide collections of accessions, very little (if any) significant associations for growth parameters were found [60], even when using very anti-conservative thresholds [2, 61] or using morphological parameters with higher heritabilities [62]. Even with more targeted growth traits like root cell length, Meijon *et al.* did not detect any significant signal above the threshold, although the first peak just below the threshold identified a causal gene [63]. Overall, it is argued that linkage and GWA studies are complementary in the loci that they are able to reveal, depending on the genetic architecture of the trait in the population considered (e.g. [21, 64]). Here, by studying four different crosses to the reference Col-0, we find many cross-specific loci, especially of mild effects, several of which might correspond to low frequency-alleles that would not likely be pictured in GWAS (not enough power due to low frequency x effect size).

Whether linkage or association mapping, these approaches are both similarly phenotyping-intensive and prone to interaction with uncontrolled environmental parameters (increasingly so with the scale of the experiments). Using our high-throughput phenotyping robots to grow individual plants under tightly controlled conditions, we intended to dissect the genetic basis of plant growth under optimal and limiting watering conditions, to a level of accuracy rarely reached so far. We focused on vegetative growth from days 8 to 29 (after sowing) and selected three non fully-correlated variables allowing to characterize plant growth dynamically: rosette area (PRA), relative growth rate (RER) and compactness. PRA is a typical cumulative trait: what is observed on day n is not independent from what has occured from day 1 to n-1.

Compactness, which basically represents a measure of PRA normalised by rosette width, is rather independent on cumulative phenomenons and hence shows much more age-specific QTLs; this is particularly striking when comparing the dynamic analyses for these traits. Finally RER integrates growth rates over a specific period of time, and is independent from plant size (= relative). Because estimated during the exponential phase of growth, RER is very much similar to what is obtained by fitting exponential models to the PRA data and exploiting model parameters [2].

Hence, growth-related phenotypes, depending on how they are exploited, will present different genetic architectures throughout age, with individual loci making different contributions to cumulative or age-specific traits. It has previously been shown that heritabilities for growth-related traits change over time [6, 65], suggesting that QTLs are more or less likely to act at specific time points. For instance, studying growth dynamics in maize [66, 67] and root tip growth in Arabidopsis [68] allowed to identify marker-trait associations that would not be detectable by considering the cumulative trait only at a single (final) time point. At the other end of time-resolution for growth, going into much more details of the dynamics (several images per day) may result in noisy raw data requiring further treatment before exploiting, for example due to projected growth estimates interfering with circadian leaf movements [65].

Our work has been performed under two environmental conditions, a control condition and one that moderately limits growth due to water (but not nutrient) availability [34], i.e. a mild drought treatment. The QTL profiles obtained at the genomic scale are very robust to mild drought with most of the large effect QTL showing no clear signs of interaction with water availability in our conditions. Some of them still change their level of significance with conditions, but it is difficult to know if this is a real interaction with drought, a change in trait variance under stress, or a change in the rest of the genetic architecture of the trait (which will impact the significance of individual QTL). Condition-specific QTLs detected here always are of small effect, which also raises the question of the power of these comparisons across different conditions/experiments due to false negative in mapping QTL. Still, this result (a relatively smaller part of the phenotypic variation is plastic rather than constitutive) is similar to what was found previously for instance in linkage [69] and association mapping [61], showing an overlap in the network of genes that regulate plant size under control and mild drought conditions [60]. Drought stress might have pleiotropic effects on different tightly interrelated phenotypic traits and impose strong constraints on them, reducing trait variability [31]. Of course, this may highly depend on the type of stress (intensity, stage of application, stability…) that is applied. Here, we chose a mild stress intensity, to remain physiologically relevant and avoid the squeezing of trait variation concomitant to strong stress levels. Also, our robot is compensating (twice a day) the individual plant size effect on transpiration that may otherwise artificially increase stress intensity according to intrinsic plant size difference.

One advantage of using a star-like cross design, increasingly used in nested association mapping (NAM) populations, is to combine the power of individual crosses and take advantage of the comparison of multiple alleles, with respect to the reference allele in order to identify allelic variants more efficiently [70–73]. These could correspond to variants originating from Col-0 or to shared allelic variants among the other parents (same direction of the allelic effect in all crosses), or to allelic series (other parents could have divergent direction/intensity of effect with respect to the reference parent). This multi-population study has confirmed the effect of several loci across traits and environments, with particular power for compactness, and in some instances already allows to predict that independent loci actually underlie major peaks. However, this approach is still limited by the mapping resolution which makes it difficult to distinguish shared variants from linked loci.

Considering the precision of our phenotyping (which has an impact on the part of phenotypic variation that is amenable to genetic dissection), the need for higher-resolution approaches to better describe the trait’s genetic architecture is obvious here. If the architecture of variation in our crosses is more complex than just a limited number of loci independently segregating for intermediate or major effects, then the density of recombination observed in a simple RIL set will not allow to decipher the full architecture [74, 75]. We first went deeper in resolution by phenotyping numerous pairs of near-isogenic lines (actually, HIFs) that each interrogate a specific portion of a chromosome (2-3Mb on average) in an otherwise fixed genetic background. This nicely confirms a majority of loci but also shows some effects that were unexpected after the initial QTL mapping results, already indicating more complex genetic architecture than anticipated; this includes single peaks splitting up in independent loci or complex patterns of linkage versus pleiotropy (when comparing different traits) and linkage versus GxE (when comparing traits in different conditions). It seems that QTL colocalisation among crosses (linkage versus shared variants) is also often questioned, although this requires to be able to compare multiple HIFs and they may not be directly comparable due to interactions with the genetic background. The difficulties to identify genes responsible for complex phenotypes also depends on their involvement in epistatic interactions. HIFs are particularly sensitive to epistasis (compared to traditional NILs) as they each harbour a different genetic background, which means that they allow epistasis to be interrogated providing that one can test enough independent HIFs, otherwise they have to be compared with care. Epistatic interactions are detected in all crosses / traits (although not always with very high significance) even when the trait heritability or variation is not so high in a cross, illustrating another factor of the complex genetic architecture. Some interactions seem to be condition-specific, but power issues are likely to be limiting in understanding these patterns of GxGxE. Furthermore, HIF have an expected candidate segregating region of several Mb usually, so our observations are likely just a glimpse at the real complexity of growth as it is known that linkage and epistasis is also active at a very local scale [44].

Still, the sensitivity of our approach here is validated by the detection of QTL colocalising with several already-known QTG expected to segregate in our crosses, like *CRY2* as discussed above [52, 76], *MPK12* which would explain nicely the bottom of chromosome 2 QTL in CvixCol [77, 78], or SG3 detected here through its epistasis with SG3i in BurxCol [49, 79].

To avoid genome-wide epistasis and better reveal local-scale architecture, we have investigated in further details a single HIF background for a specific 3Mb region containing a known QTG of large and pleiotropic effect (*CRY2*) in an original approach. Our analysis reveals that there are at least 3 other QTGs in this interval controlling one of the traits in at least one condition. For PRA29, the picture seems to be even more complex with traces of at least one more locus; here, it seems that phenotyping accuracy becomes again limiting after all. Obviously these loci with opposite allelic effects, different patterns of pleiotropy and interaction with the environment, and just a few hundreds of kb from each other, remain cryptic in simple QTL mapping. This major result of local-scale independent complex genetic architecture for different traits and conditions should lead us to a lot of caution when interpreting colocalising QTLs from different traits / conditions / age, as these may very well be independent loci rather than a single pleiotropic locus, as shown here for PRA and Compactness. If we were to extrapolate the figures obtained on this specific region in this particular cross to the genome- and species-scale, we would expect hundreds of causative loci of detectable phenotypic effect controlling these growth-related phenotypes. One way to approach these individual loci would be to decompose their independent signature based on different dynamics or underlying traits (transcriptomics, metabolomics…) in a ‘systems genetics’ strategy [54, 60, 80].

Complex genetic architecture as revealed in this study has consequences on quantitative genetics experimental design and interpretation, arguing in favor of linkage mapping or GWAS depending on the balance between genetic complexity due to linked loci (where association is expected to behave better than linkage mapping) and genetic complexity due to small effect/rare alleles (where association will behave poorly). Other intermediate experimental designs like multiparental populations or nested-association mapping should bring more power [12, 81]. Resolution is improved by pushing recombination densities to its limits and it was shown to help resolve more complex genetic architecture in yeast [82]. In plants, using ‘hyper-recombinant’ mutations to generate new segregating populations could also be a strategy in the future [83].

## Materials And Methods

**Genetic material. Recombinant Inbred Lines (RILs):** The 4 RIL sets used for this work were generated at the Versailles Arabidopsis Stock Center, France (http://publiclines.versailles.inra.fr/) and described previously [47]. Versailles stock center ID are indicated as ‘xxxAV’ and ‘xxRV’ for Accessions and RILs respectively). They are derived from crosses between the following pairs of accessions :

RIL set ‘BlaxCol’ (ID = 2RV): Bla-1 (76AV) x Col-0 (186AV) / 259 RILs

RIL set ‘CvixCol’ (ID = 8RV): Cvi-0 (166AV) x Col-0 / 358 RILs

RIL set ‘BurxCol’ (ID = 20RV): Bur-0 (172AV) x Col-0 / 283 RILs

RIL set ‘YoxCol’ (ID = 23RV): Yo-0 (250AV) x Col-0 / 358 RILs

**Heterogeneous Inbred Families (HIFs):** HIFs have been selected in each RIL set to cover several regions of the genome, some of which are expected to segregate for QTLs while others were chosen at random locations. As described previously [84] they are derived from the progeny of one RIL which is heterozygous only at the locus of interest. Hence, one HIF family is composed of 2 or 3 lines fixed for one parental allele at the segregating region and 2 or 3 lines fixed for the alternate parental allele at the segregating region, in an otherwise identical genetic background. Each HIF (ID = ‘xxHVyyy’) is named after the RIL ID (‘xxRVyyy’) from which it has been generated (‘xx’ is the ID of the RIL set, ‘yyy’ is the ID of the RIL): for example the family 2HV142 is fixed from the RIL 2RV142. The tentative QTL validation is based on the phenotypic comparison of these fixed lines within the HIF family. The complete dataset gathering genotypic information on the RILs used to generate HIFs has been submitted to INRA institutional data repository (https://data.inra.fr) [50], with the genotypic conventions and ID from the Versailles Arabidopsis Stock Center (http://publiclines.versailles.inra.fr/).

**microStairs:** The progeny of CvixCol RIL 8RV294 (also used to generate HIF family 8HV294), segregating for the first 3Mb of chromosome 1, has been screened to detect recombined individuals. 4000 individuals were genotyped with markers at the edge of the heterozygous region and then 29 evenly distributed recombinants were selected using markers spaced every ˜100kb. These recombinants were genotyped and fixed for the remaining segregating region in such a way that they each differ genotypically from the next recombinant by a ˜100kb bin on average (@Suppl. Table S4). This is similar to the approach taken by Koumproglou *et al.* [85], but at a much finer scale, hence the name ‘*microStairs*’.

**Phenotyping:** Phenotyping was performed on the Phenoscope robots as previously described [34] (https://phenoscope.versailles.inra.fr/). Every RIL set and their respective parental accessions have been phenotyped in 2 independent Phenoscope experiments (= biological replicates), except for CvixCol (3 biological replicates). In short, the peatmoss plugs’ soil water content (SWC) is gradually adjusted for each plant individually as a fraction of the initially-saturated plug weight. We worked at 2 watering conditions: 60% SWC for non-limiting conditions (called ‘WW’ for well watered) and 30% SWC for mildly growth-limiting watering conditions (called ‘WD’ for water deficit). The growth room is set at a 8 hours short-days photoperiod (230 µmol m-2 sec-1) with days at 21°C/65%RH and nights at 17°C/65%RH. A picture of each individual plant is taken every day at the same day-time and a semi-automatic segmentation process (with some manual corrections when required) is performed to extract leaf pixels. From this we exploit different traits: Projected Rosette Area (PRA), circle radius, convex hull area, average Red, Green and Blue components (leaf pixels, RGB colour scale), and derived phenotypic traits are calculated, such as the compactness (ratio PRA / convex hull area) and the Relative Expansion Rate (RER) over specific time windows (@Figure 1), as previously described [34]. The complete raw phenotypic dataset has been submitted to INRA institutional data repository (https://data.inra.fr) [48]. Principal component analysis (PCA) was performed using the ade4 R package based on phenotypic data from all the RIL sets in WW and WD conditions. For QTL detection and further analyses, the phenotypic values were corrected for experiment effects. Corrected phenotypic values were calculated using the intercept (μ), the condition (α), genotype (γ), and genotype*condition (σ) effects of the following model :

Y_ijkl ˜ μ + α_i + β_j + γ_k + δ_ij + λ_jk + σ_ik + ε_ijkl

where Y_ijkl: phenotype; μ: mean; α_i: effect of the condition; β_j: effect of the experiment; γ_k :effect of the genotype; δ_ij :effect of the interaction condition*experiment; λ_jk :effect of the interaction experiment*genotype; σ_ik :effect of the interaction condition*genotype; ε_ijkl: residuals.

Note that one of the replicates of the CvixCol phenotypic data was already analysed for QTL mapping -with a different statistical model- in Tisné *et al.* [34].

Similarly, the set of near isogenic lines (microStairs) were phenotyped and analysed from 3 full biological replicates in independent Phenoscope experiments.

**QTL MQM:** QTL detections were performed using Multiple QTL Mapping algorithm (MQM) implemented in the R/qtl package [86, 87] using a backward selection of cofactors. At first, genotype missing data were augmented, then one marker every three markers were selected and used as cofactors. Important markers were selected through backward elimination. Finally, a QTL was moved along the genome using these pre-selected markers as cofactors, except for the markers in the 25.0 cM window around the region of interest. QTL were identified based on the most informative model through maximum likelihood. According to permutation tests results, a universal LOD threshold of 2.4 was chosen for all QTL maps. Interactive QTL maps for time-course series were generated using the R/qtl charts package [88]. All QTL positions were projected on the consensus genetic map of the 4 crosses built with R/qtl. A joint genotype dataset was constructed with ‘A’ alleles coding Col alleles (the common parent), ‘B’ alleles for non-Col alleles, and monomorphic markers in a cross coded as missing. The linkage groups were considered known from the individual maps and the physical position of markers, and a first marker order was calculated using orderMarkers function. In case of conflicting marker order between individual, physical and consensus maps, the function switch.order was used to retain the most probable order (i.e with the lowest number of recombination).

**Epistatic interactions:** Epistatic interactions were identified using the scantwo function of the R/qtl package. LOD scores were calculated for additive, interaction and full models. LOD score and p-values were calculated for all pairwise combination of markers, except for adjacent markers. Effect plots for the pairs of markers were drawn using the R/qtl package.

**QTL mapping in the multi-cross design:** QTL mapping in the multi-cross design was performed with the MCQTL package [89]. The model was described as additive (no dominance effect) and connected (Col-0 centered design) and the following 3 steps process was applied. Step 1: thresholds were calculated by variable on the whole genome using 1000 resampling replicates (PRA29_WW = 3.73; PRA29_WD = 3.84; RER16-29_WW = 5.22; RER16-29_WD = 3.95; Compactness29_WW = 3.64 Compactness29_WD = 3.30). Step 2: QTL detection was performed using iQTLm method with a threshold of 4. To perform this detection, cofactors were automatically chosen by backward selection with a threshold of 2.8 among a skeleton with a minimal inter distance of 10cM. Search for QTL was not allowed within +/-10cM window surrounding the QTL to avoid linked genetic regions. Step 3: model estimations were performed for each variable and condition using the parameters previously described and the QTL position identified at the QTL detection step.

**Fine dissection of a genomic region ‘microStairs’:** The phenotypes of the recombined HIFs lines were modeled using the following linear equation :

Y_ij ˜ μ + α_i + β_j + ε_ij

where Y_ij is the value of the phenotype; μ is the mean of the phenotype; α_i is the effect of the stair (bin) i; β_j is the effect of the line j and ε_ij is the residuals. An anova was performed with this model and the p-value of the stair effects were adjusted by a Benjamini-Hochberg correction.

Polymorphic candidate genes (Cvi versus Col) were listed for each PRA29 ‘microStairs’ significant interval according to variants listed on the 1001Genomes website (http://1001genomes.org/), through the Polymorph1001 tool. Differentially cis-regulated variants were extracted from Cvi/Col Allele-Specific Expression (ASE) data [57] and from CvixCol local-eQTLs data [90] across the whole region.

## Acknowledgments

We thank Yann Serrand for the supervision of the *Phenoscope*, we thank Francisco Cubillos, Cécile Grondin, Isabelle Gy and Mathieu Canut for the selection of recombinants from 8HV294. We also thank Laurence Moreau for her help in performing multi-cross analyses. We thank José Jiménez-Gómez for discussions on statistical analyses of the ‘microStairs’ experiment. This work was supported by funding from the European Commission Framework Programme 7, ERC Starting Grant ‘DECODE’/ERC-2009-StG-243359 to O.L. and Agence Nationale de la Recherche (ANR) grant ‘2Complex’/ANR-09-BLAN-0366 to O.L. The IJPB benefits from the support of the LabEx Saclay Plant Sciences-SPS (ANR-10-LABX-0040-SPS).

## Datasets

> Two datasets have been submitted to INRA Dataverse repository (https://data.inra.fr):

### - Raw phenotypic data obtained on the Arabidopsis RILs with the Phenoscope robots [48]

This dataset gathers the main raw phenotypic data obtained and exploited in Marchadier, Hanemian, Tisné *et al.* (2018). It contains data from 4 RIL sets across 9 Phenoscope experiments.

For each Phenoscope experiment, Recombinant Inbred Line (RIL) and Condition (‘WW’ = Well Watered / ‘WD’ = Water Deficit), the data set indicates the phenotypic value for 6 traits at 21 successive time points. ‘Trait.XX’ = Trait at XX days after sowing, with ‘XX’ = 09 to 29 and ‘Trait’ = PRA (Projected Rosette Area; in cm2), GreenMean / RedMean / BlueMean (rosette pixels’ colour components; arbitrary unit), ConvexHullArea (area of the convex hull encompassing the rosette; in cm2) and CircleRadius (radius of the smallest circle encompassing the rosette; in cm). RIL set IDs and RIL IDs are according to Publiclines http://publiclines.versailles.inra.fr/rils/index

### Genotypic description of the near isogenic lines (HIFs) used for QTL validation and significance of the observed segregating phenotypes [50]

Each row represents a single HIF and the genotype of the F7 RIL it originates from is indicated along the chromosomes with RIL ID, markers and genotypic conventions from Publiclines http://publiclines.versailles.inra.fr/rils/index (i.e. ‘A’ = Col allele; ‘B’ = alternate parental allele; ‘C’ = heterozygous). The region highlighted in yellow is the segregating region that is tested in the HIF family through several fixed lines for each parental allele. For each of the 3 growth traits (Compactness29 = rosette compactness 29 days after sowing; PRA29 = Projected Rosette Area 29 days after sowing; RER16-29 = Relative rosette Expansion Rate between days 16 and 29 after sowing) in 2 conditions (‘WW’ = Well Watered / ‘WD’ = Water Deficit), whenever significant, the p-value of the comparison between allelic lines (‘Pval’) and the direction of the allelic effect (‘sign’ calculated as [Col-Xxx] where Xxx is the alternate parental allele) are indicated in the last columns of the table.

> A specific webpage is associated with this work to display interactive graphes for dynamic QTL analyses at http://www.inra.fr/vast/PhenoDynamic.htm

## Supplementary Material

**Supplementary Figure S1:**
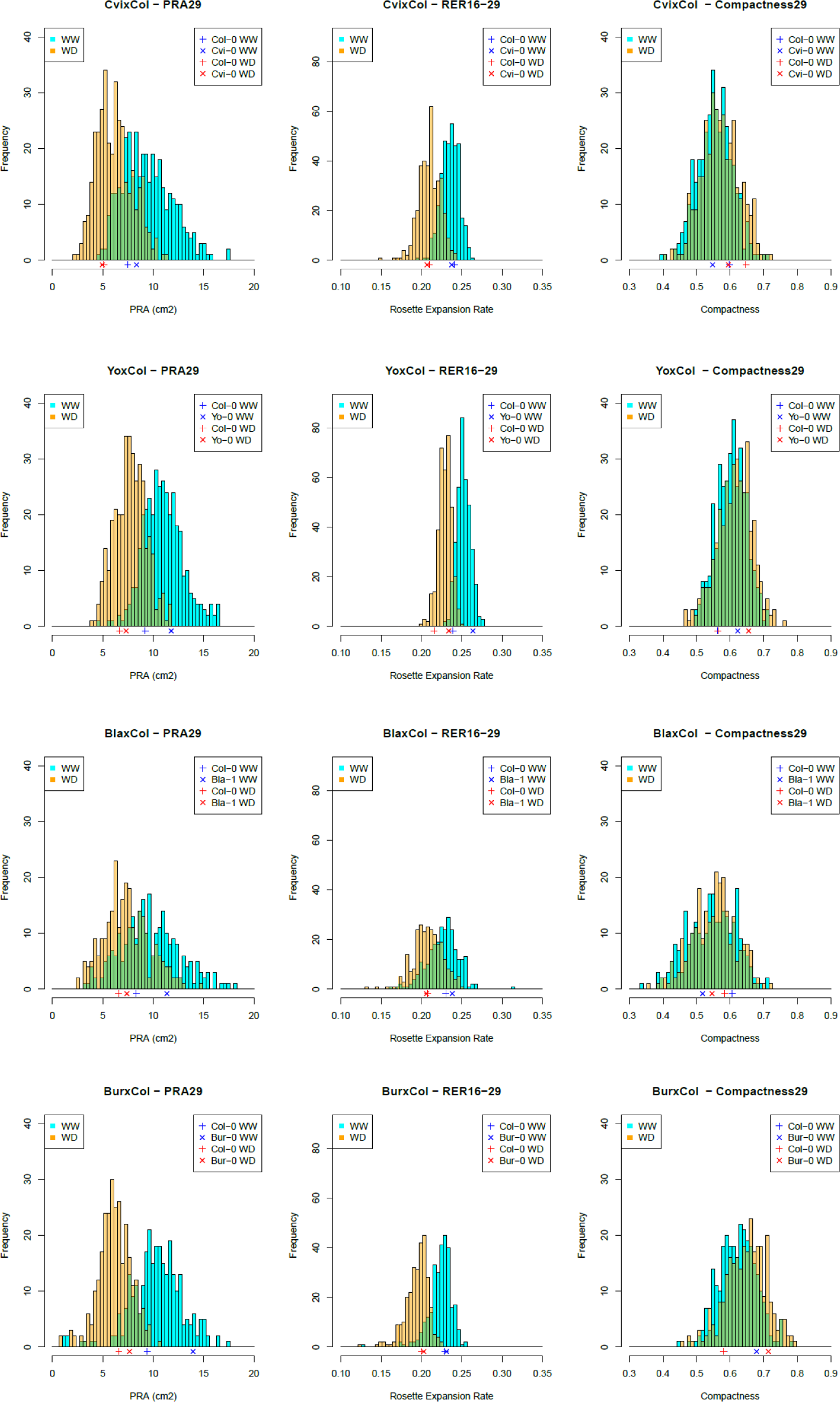
Distributions of the mean phenotypic values across RILs. Frequency histograms showing the distributions of the PRA29, RER16-29 and Compactness29 traits within each of the four RIL sets in well watered (WW: blue) and water deficit (WD: orange) conditions. Phenotypic values for parental accessions from these specific experiments are indicated by blue (WW) and red (WD) ticks (see inset for legend) just above the x axes.

**Supplementary Figure S2:**
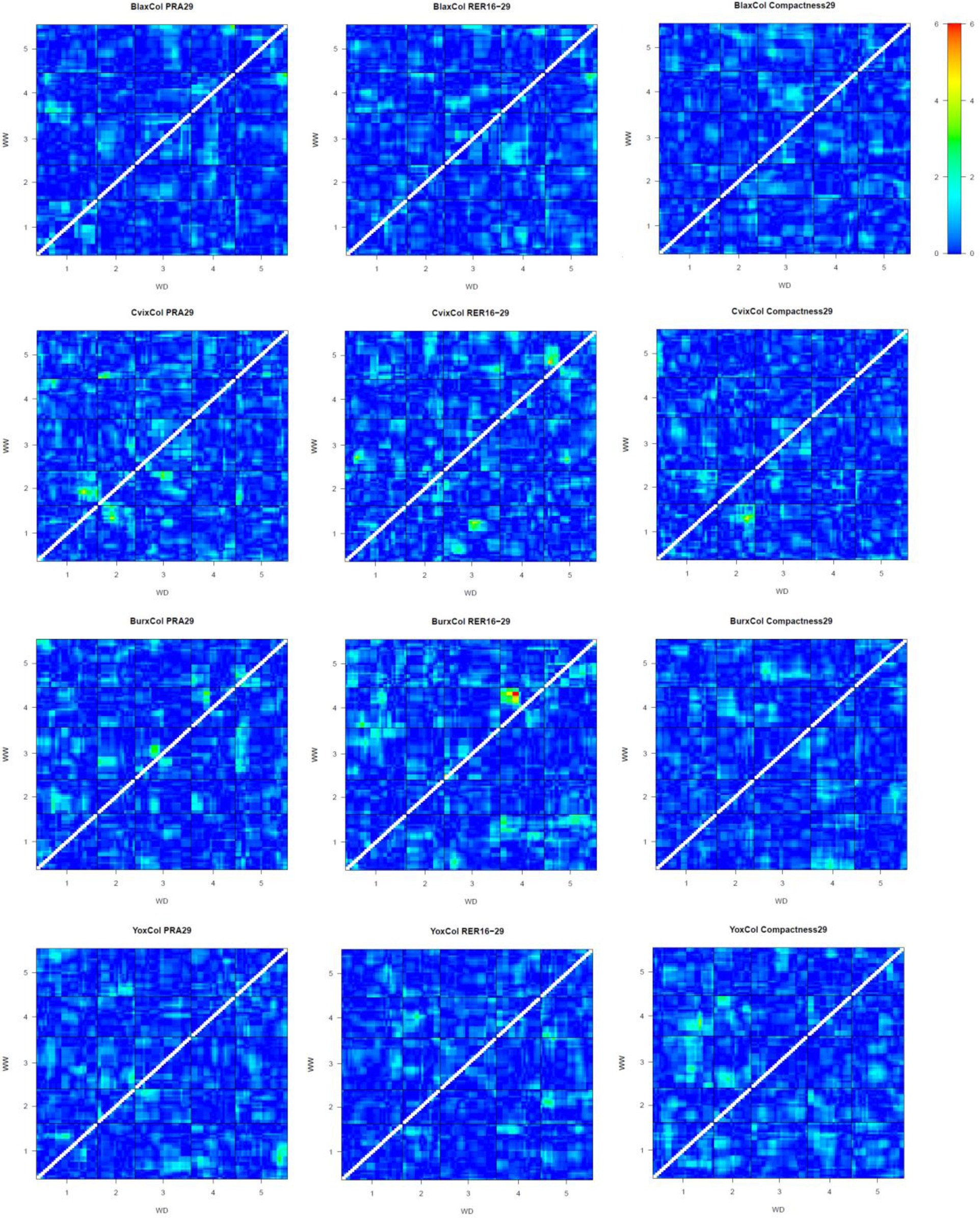
2D scans for epistasis. Heatmaps representing the LOD score of the interaction effects between all pairs of loci for 4 RIL sets, 3 traits, in 2 conditions. Each heatmap shows the pairwise interaction effects obtained in control condition WW (triangle above diagonal) and in water deficit condition WD (triangle below diagonal). The color scale (LOD score values) shared among all heatmaps is indicated on the right (note that it is different from the scale of @Figure 4). Diagonal values are canceled and enlarged to exclude pairs of adjacent markers from the test.

**Supplementary Figure S3:**
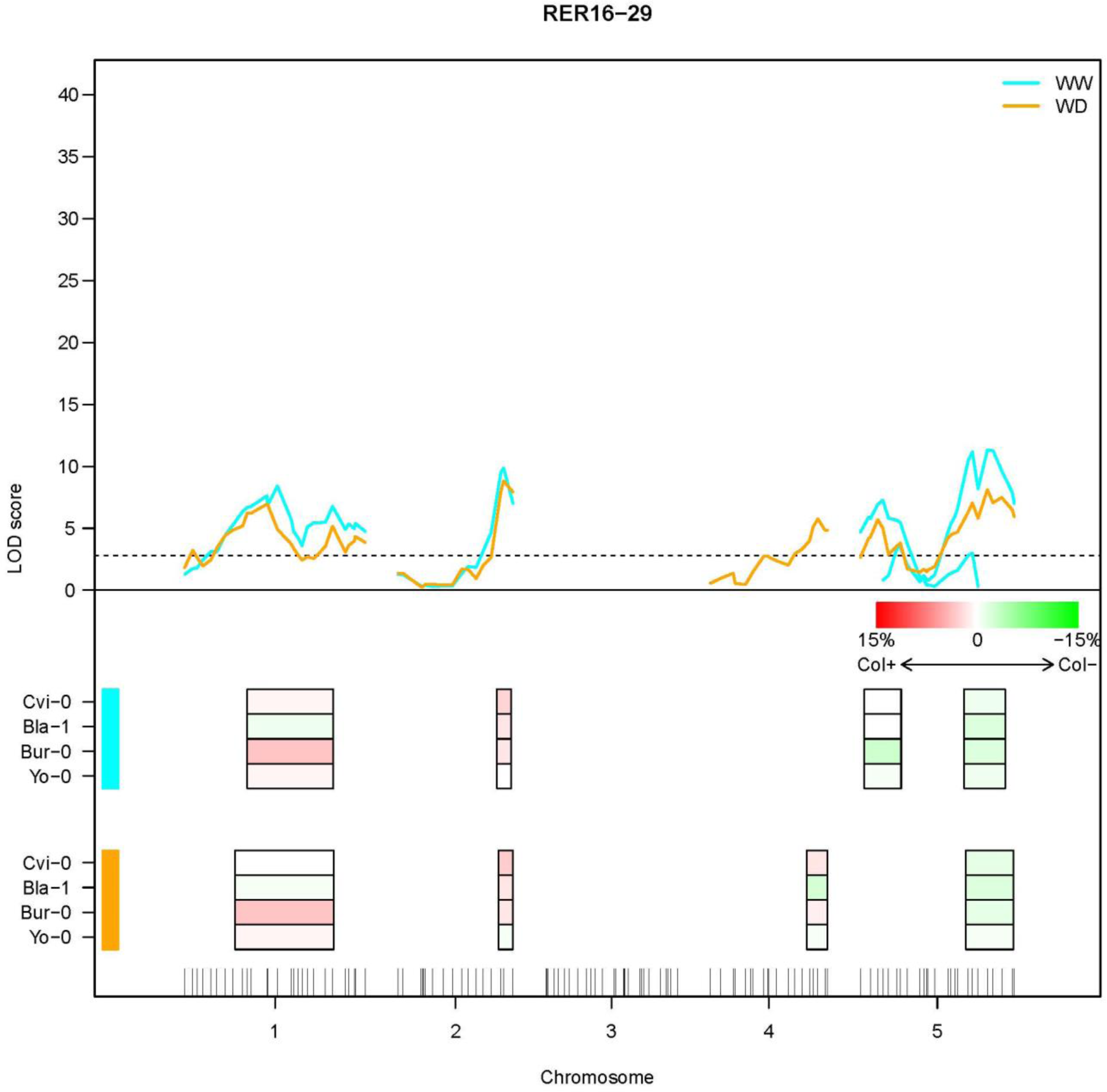
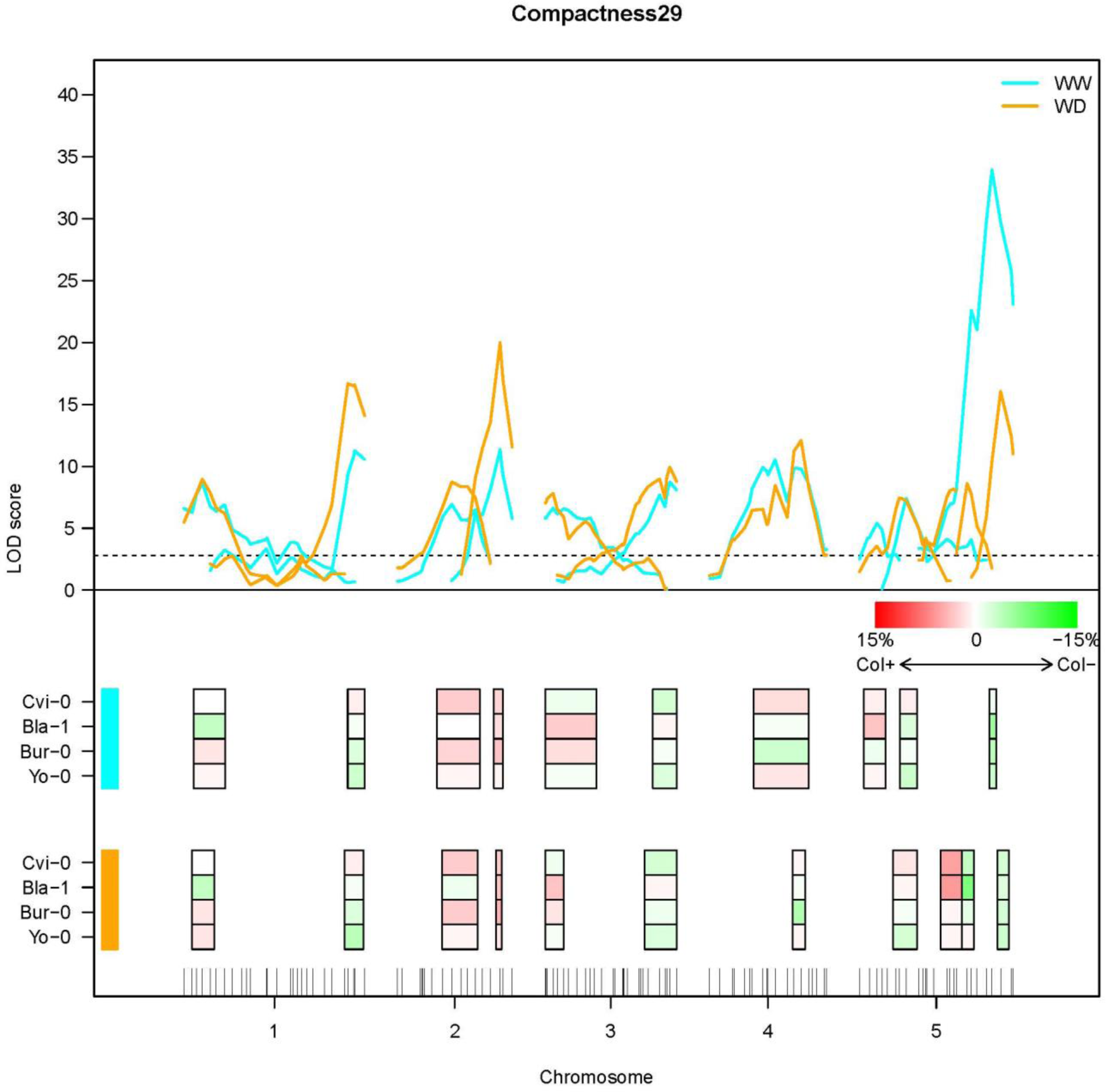
Multi-cross QTL analysis for RER16-29 and Compactness29. Same legend as @Figure 5. Chromosome 3 shows no significant combined QTLs for RER16-29.

**Supplementary Figure S4:**
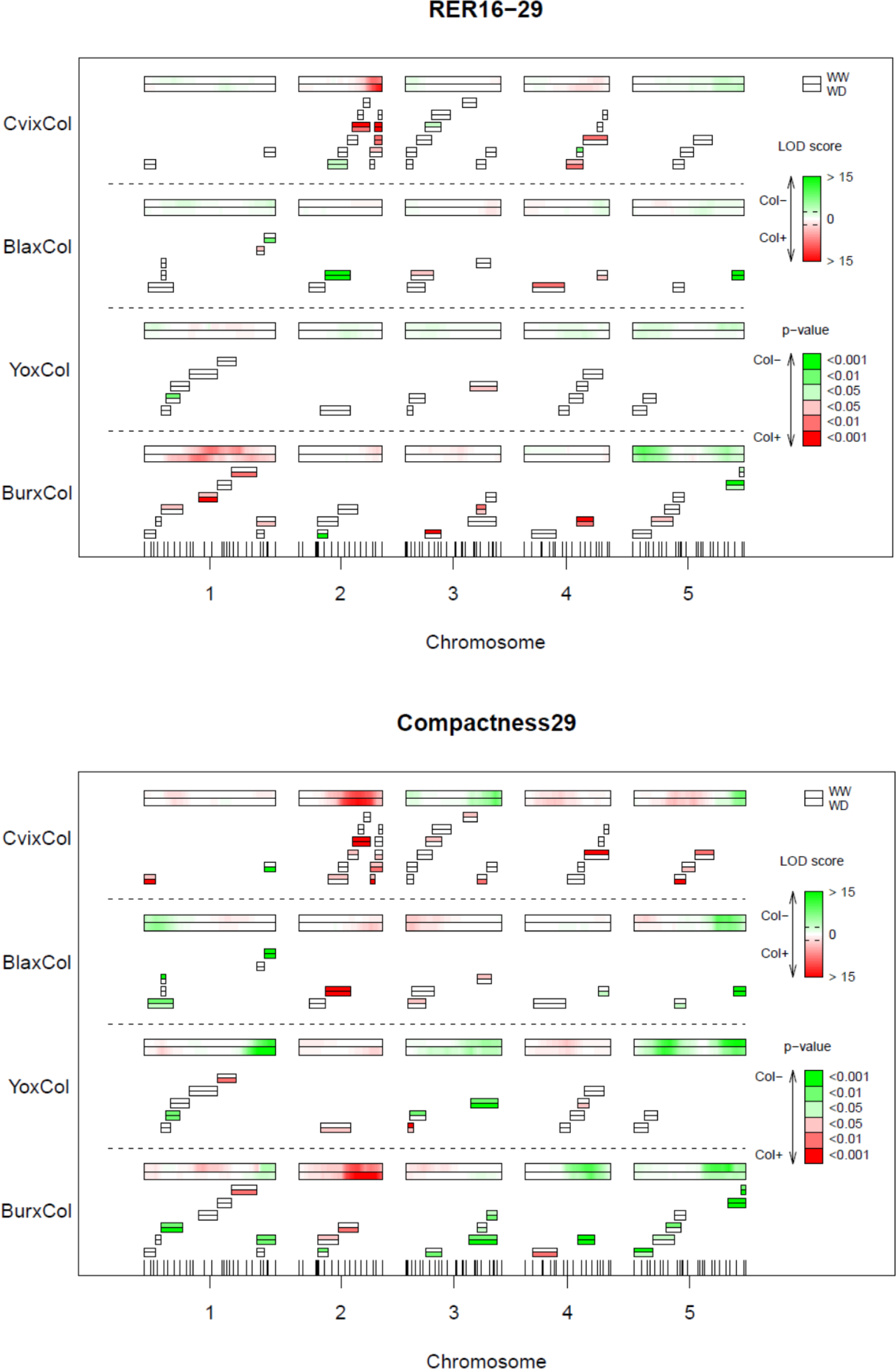
Near isogenic lines-based validation of QTLs for RER16-29 and Compactness29. Same legend as @Figure 6

**Supplementary Figure S5:**
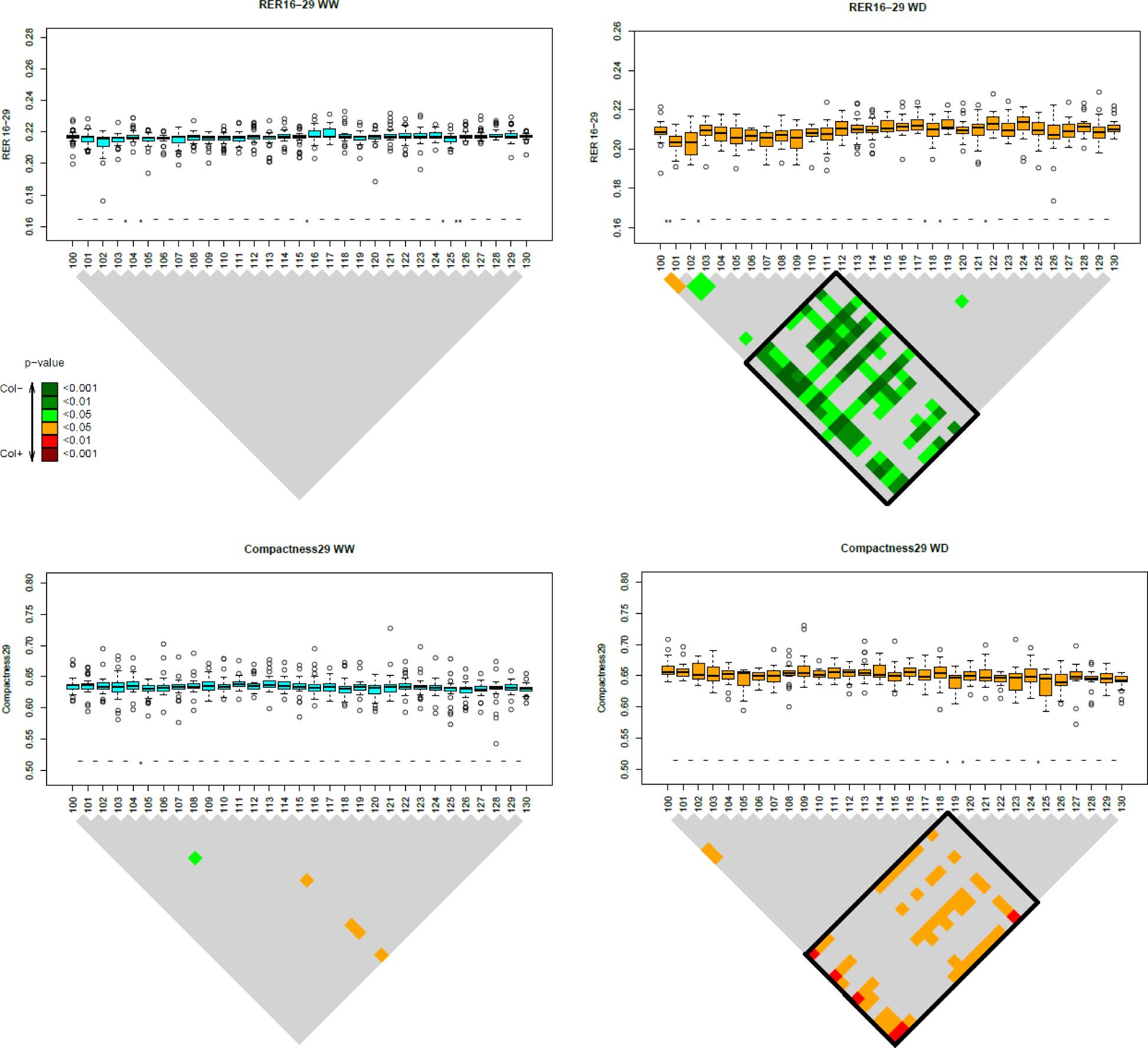
Dissection of a genomic region in CvixCol for RER16-29 and Compactness29 (‘microStairs’ approach) Same legend as @Figure 7

**Supplementary Table S1:** Heritabilities of the observed phenotypes. Broad-sense heritabilities (h2) in the 4 RIL sets (BlaxCol, CvixCol, BurxCol and YoxCol) for PRA29, RER16-29 and Compactness29 in WW and WD conditions. H2 > 0.4 are highlighted in bold. Broad-sense heritabilities were calculated with the following equation h2 = Var(G)/Var(P) with Var(P)=Var(G)+Var(E)

|  | BlaxCol | CvixCol | BurxCol | YoxCol |
| --- | --- | --- | --- | --- |
| PRA29 WW | 0.23 | **0.67** | **0.62** | **0.48** |
| PRA29 WD | 0.12 | **0.45** | **0.55** | 0.26 |
| RER16-29 WW | 0.17 | 0.13 | **0.66** | NA |
| RER16-29 WD | 0.22 | NA | **0.42** | NA |
| Compactness29 WW | **0.75** | **0.60** | **0.65** | **0.54** |
| Compactness29 WD | **0.50** | **0.40** | 0.37 | 0.34 |

**Supplementary Table S2:** Analyses of variance for genotype, experiment and condition factors, and their interactions. For each RIL set, -log(p values) of main and interaction effects calculated by the analysis of variance (ANOVA) from the following model : Yijkl˜μ+αi+βj+γk+δij+λjk+σik+εijkl where Yijkl: phenotype; μ: mean; αi: effect of the condition; βj: effect of the experiment; γk: effect of the genotype; δij: effect of the interaction condition*experiment; λjk: effect of thefig interaction experiment*genotype; σik: effect of the interaction condition*genotype; εijkl: residuals NS = Not Significant

| RIL set | Phenotype | Condition | Experiment | Genotype | Condition * Experiment | Experiment * Genotype | Condition * Genotype |
| --- | --- | --- | --- | --- | --- | --- | --- |
| CvixCol | PRA29 | 726,57 | 26,33 | 526,59 | 75,81 | 51,37 | 16,87 |
| CvixCol | Compactness29 | 57,54 | 212,95 | 378,58 | 50,64 | 13,01 | NS |
| CvixCol | RER16-29 | 555,13 | 499,28 | 138,67 | 69,48 | 8,13 | 3,06 |
| YoxCol | PRA29 | 450,11 | 27,20 | 185,91 | 20,90 | 47,28 | 3,30 |
| YoxCol | Compactness29 | 12,03 | 96,39 | 151,05 | 25,42 | NS | NS |
| YoxCol | RER16-29 | 482,02 | 355,28 | 84,52 | 8,39 | 6,57 | NS |
| BlaxCol | PRA29 | 211,83 | 10,96 | 212,99 | NS | 184,35 | 9,44 |
| BlaxCol | Compactness29 | NS | 4,27 | 183,33 | NS | 4,85 | 4,96 |
| BlaxCol | RER16-29 | 161,24 | 47,86 | 112,12 | NS | 73,21 | 3,69 |
| BurxCol | PRA29 | 402,90 | 14,73 | 168,74 | 7,37 | 4,92 | 11,24 |
| BurxCol | Compactness29 | 34,46 | 8,21 | 91,88 | NS | NS | NS |
| BurxCol | RER16-29 | 407,91 | 22,25 | 149,56 | NS | 11,60 | 13,24 |

**Supplementary Table S3:** Mapped QTL parameters across RIL sets, traits and conditions. For each RIL set, trait and condition (or GxE interaction term: ‘inter’): every independent QTL peak is a row in the table, indicating its localisation (chromosome and position on the genetic map), its significance (maximum LOD score) and its effect (direction of the allelic effect: ‘sign’ indicates the sign of the allelic effect estimated as [Col – Xxx], where Xxx is the alternate parental allele; R2 %). Independent QTLs peaks are considered when the LOD score curve returns below threshold between 2 peaks, to ensure the loci are not too genetically linked.

| RIL set | Phenotype | Condition | Chromosome | Position (cM) | LOD score | sign | R2 (%) |
| --- | --- | --- | --- | --- | --- | --- | --- |
| BurxCol | PRA29 | WD | 1 | 5 | 4,1 | -1 | 1 |
| BurxCol | PRA29 | WD | 1 | 50 | 11,2 | 1 | 11 |
| BurxCol | PRA29 | WD | 5 | 93 | 2,6 | -1 | 3 |
| BurxCol | PRA29 | WW | 1 | 11 | 9,1 | -1 | 4 |
| BurxCol | PRA29 | WW | 1 | 50 | 19,8 | 1 | 12 |
| BurxCol | PRA29 | WW | 4 | 0 | 8,3 | -1 | 9 |
| BurxCol | PRA29 | WW | 5 | 14 | 2,8 | -1 | 2 |
| BurxCol | PRA29 | WW | 5 | 93 | 3,2 | -1 | 3 |
| BurxCol | PRA29 | inter | 1 | 20 | 9 | -1 | 7 |
| BurxCol | PRA29 | inter | 1 | 50 | 6,6 | 1 | 4 |
| BurxCol | PRA29 | inter | 2 | 70 | 3,9 | 1 | 3 |
| BurxCol | PRA29 | inter | 4 | 0 | 5,3 | -1 | 6 |
| BurxCol | RER16-29 | WD | 1 | 50 | 9,3 | 1 | 11 |
| BurxCol | RER16-29 | WD | 1 | 74 | 3,1 | 1 | 4 |
| BurxCol | RER16-29 | WD | 1 | 90 | 2,9 | 1 | 8 |
| BurxCol | RER16-29 | WD | 5 | 11 | 9,3 | -1 | 6 |
| BurxCol | RER16-29 | WD | 5 | 86 | 4,3 | -1 | 3 |
| BurxCol | RER16-29 | WW | 1 | 56 | 12,4 | 1 | 9 |
| BurxCol | RER16-29 | WW | 2 | 64 | 2,9 | 1 | 2 |
| BurxCol | RER16-29 | WW | 5 | 11 | 16 | -1 | 12 |
| BurxCol | RER16-29 | WW | 5 | 80 | 8 | -1 | 4 |
| BurxCol | RER16-29 | inter | 1 | 25 | 3,2 | -1 | 5 |
| BurxCol | Compactness29 | WD | 1 | 110 | 4,1 | -1 | 3 |
| BurxCol | Compactness29 | WD | 2 | 56 | 16,4 | 1 | 11 |
| BurxCol | Compactness29 | WD | 4 | 56 | 12 | -1 | 8 |
| BurxCol | Compactness29 | WD | 5 | 86 | 7,4 | -1 | 3 |
| BurxCol | Compactness29 | WW | 1 | 104 | 7,2 | -1 | 3 |
| BurxCol | Compactness29 | WW | 1 | 50 | 2,8 | 1 | 1 |
| BurxCol | Compactness29 | WW | 1 | 65 | 2,9 | 1 | 1 |
| BurxCol | Compactness29 | WW | 2 | 47 | 14,6 | 1 | 11 |
| BurxCol | Compactness29 | WW | 3 | 14 | 3,7 | 1 | 3 |
| BurxCol | Compactness29 | WW | 4 | 56 | 13,7 | -1 | 6 |
| BurxCol | Compactness29 | WW | 5 | 6 | 4 | -1 | 1 |
| BurxCol | Compactness29 | WW | 5 | 80 | 17 | -1 | 8 |
| BurxCol | Compactness29 | inter | 3 | 59 | 2,7 | 1 | 4 |
| YoxCol | PRA29 | WD | 1 | 20 | 4 | -1 | 4 |
| YoxCol | PRA29 | WD | 1 | 56 | 3,6 | -1 | 5 |
| YoxCol | PRA29 | WD | 2 | 33 | 4,2 | -1 | 3 |
| YoxCol | PRA29 | WD | 4 | 21 | 2,7 | -1 | 3 |
| YoxCol | PRA29 | WD | 4 | 47 | 2,9 | -1 | 4 |
| YoxCol | PRA29 | WD | 4 | 60 | 2,5 | -1 | 4 |
| YoxCol | PRA29 | WD | 5 | 80 | 11,6 | -1 | 9 |
| YoxCol | PRA29 | WD | 5 | 6 | 3,9 | 1 | 3 |
| YoxCol | PRA29 | WW | 1 | 16 | 3,1 | -1 | 4 |
| YoxCol | PRA29 | WW | 1 | 35 | 5,7 | -1 | 7 |
| YoxCol | PRA29 | WW | 2 | 39 | 3 | -1 | 4 |
| YoxCol | PRA29 | WW | 4 | 15 | 6,2 | -1 | 4 |
| YoxCol | PRA29 | WW | 5 | 77 | 16,8 | -1 | 12 |
| YoxCol | PRA29 | WW | 5 | 11 | 9,6 | 1 | 6 |
| YoxCol | PRA29 | inter | 4 | 0 | 3,6 | -1 | 3 |
| YoxCol | PRA29 | inter | 4 | 71 | 2,6 | 1 | 3 |
| YoxCol | PRA29 | inter | 5 | 68 | 5,6 | -1 | 4 |
| YoxCol | PRA29 | inter | 5 | 11 | 4,9 | 1 | 4 |
| YoxCol | RER16-29 | WD | 4 | 35 | 3,6 | -1 | 4 |
| YoxCol | RER16-29 | WD | 4 | 51 | 2,9 | -1 | 5 |
| YoxCol | RER16-29 | WD | 5 | 6 | 3 | -1 | 4 |
| YoxCol | RER16-29 | WD | 5 | 65 | 5,3 | -1 | 5 |
| YoxCol | RER16-29 | WW | 1 | 11 | 3,6 | -1 | 3 |
| YoxCol | RER16-29 | WW | 3 | 19 | 2,6 | -1 | 4 |
| YoxCol | RER16-29 | WW | 5 | 14 | 3,2 | -1 | 4 |
| YoxCol | RER16-29 | WW | 5 | 71 | 9,7 | -1 | 11 |
| YoxCol | RER16-29 | inter | 4 | 65 | 4,9 | 1 | 6 |
| YoxCol | RER16-29 | inter | 5 | 92 | 3 | -1 | 3 |
| YoxCol | Compactness29 | WD | 1 | 104 | 27,6 | -1 | 18 |
| YoxCol | Compactness29 | WD | 1 | 11 | 2,4 | 1 | 5 |
| YoxCol | Compactness29 | WD | 2 | 64 | 4,9 | 1 | 5 |
| YoxCol | Compactness29 | WD | 3 | 69 | 6,6 | -1 | 5 |
| YoxCol | Compactness29 | WD | 4 | 32 | 4,7 | 1 | 3 |
| YoxCol | Compactness29 | WD | 5 | 24 | 18,2 | -1 | 10 |
| YoxCol | Compactness29 | WD | 5 | 86 | 15,8 | -1 | 13 |
| YoxCol | Compactness29 | WW | 1 | 35 | 2,8 | -1 | 1 |
| YoxCol | Compactness29 | WW | 1 | 104 | 14,6 | -1 | 10 |
| YoxCol | Compactness29 | WW | 1 | 11 | 2,8 | 1 | 3 |
| YoxCol | Compactness29 | WW | 3 | 69 | 8,2 | -1 | 7 |
| YoxCol | Compactness29 | WW | 4 | 35 | 6 | 1 | 3 |
| YoxCol | Compactness29 | WW | 4 | 71 | 3,3 | 1 | 3 |
| YoxCol | Compactness29 | WW | 5 | 28 | 16,8 | -1 | 10 |
| YoxCol | Compactness29 | WW | 5 | 86 | 24,7 | -1 | 19 |
| YoxCol | Compactness29 | inter | 1 | 85 | 5,1 | 1 | 7 |
| YoxCol | Compactness29 | inter | 2 | 64 | 2,6 | -1 | 3 |
| BlaxCol | PRA29 | WD | 1 | 20 | 3,8 | -1 | 6 |
| BlaxCol | PRA29 | WW | 1 | 16 | 4,5 | -1 | 7 |
| BlaxCol | PRA29 | WW | 1 | 110 | 5,8 | -1 | 6 |
| BlaxCol | PRA29 | inter | 1 | 110 | 3,5 | -1 | 6 |
| BlaxCol | RER16-29 | WD | 3 | 69 | 2,8 | 1 | 4 |
| BlaxCol | RER16-29 | WD | 4 | 65 | 2,5 | -1 | 4 |
| BlaxCol | RER16-29 | WW | 1 | 104 | 2,7 | -1 | 3 |
| BlaxCol | RER16-29 | WW | 3 | 76 | 3,5 | 1 | 4 |
| BlaxCol | RER16-29 | WW | 5 | 65 | 2,5 | -1 | 5 |
| BlaxCol | RER16-29 | WW | 5 | 80 | 3,2 | -1 | 4 |
| BlaxCol | Compactness29 | WD | 1 | 11 | 6,8 | -1 | 8 |
| BlaxCol | Compactness29 | WD | 2 | 64 | 8,2 | 1 | 6 |
| BlaxCol | Compactness29 | WD | 3 | 0 | 9,6 | 1 | 9 |
| BlaxCol | Compactness29 | WD | 5 | 77 | 12,2 | -1 | 10 |
| BlaxCol | Compactness29 | WW | 1 | 11 | 6,3 | -1 | 8 |
| BlaxCol | Compactness29 | WW | 1 | 74 | 2,7 | 1 | 5 |
| BlaxCol | Compactness29 | WW | 2 | 64 | 3,8 | 1 | 2 |
| BlaxCol | Compactness29 | WW | 3 | 0 | 5,5 | 1 | 4 |
| BlaxCol | Compactness29 | WW | 5 | 77 | 22,4 | -1 | 20 |
| BlaxCol | Compactness29 | WW | 5 | 11 | 5,6 | 1 | 4 |
| CvixCol | PRA29 | WD | 1 | 20 | 4,5 | -1 | 4 |
| CvixCol | PRA29 | WD | 2 | 70 | 29,6 | 1 | 30 |
| CvixCol | PRA29 | WD | 4 | 47 | 2,6 | -1 | 1 |
| CvixCol | PRA29 | WD | 5 | 86 | 6,3 | -1 | 4 |
| CvixCol | PRA29 | WW | 1 | 16 | 9,4 | -1 | 7 |
| CvixCol | PRA29 | WW | 1 | 104 | 3 | -1 | 4 |
| CvixCol | PRA29 | WW | 2 | 64 | 27,3 | 1 | 24 |
| CvixCol | PRA29 | WW | 4 | 47 | 6,9 | -1 | 3 |
| CvixCol | PRA29 | WW | 5 | 86 | 9,4 | -1 | 4 |
| CvixCol | PRA29 | inter | 1 | 0 | 7,5 | -1 | 8 |
| CvixCol | PRA29 | inter | 2 | 42 | 3,4 | 1 | 4 |
| CvixCol | PRA29 | inter | 4 | 47 | 4,9 | -1 | 4 |
| CvixCol | RER16-29 | WD | 2 | 70 | 32 | 1 | 27 |
| CvixCol | RER16-29 | WD | 5 | 71 | 7 | -1 | 4 |
| CvixCol | RER16-29 | WW | 2 | 64 | 23,3 | 1 | 22 |
| CvixCol | RER16-29 | WW | 5 | 80 | 4,8 | -1 | 4 |
| CvixCol | RER16-29 | inter | 1 | 5 | 2,7 | -1 | 3 |
| CvixCol | Compactness29 | WD | 2 | 47 | 14,2 | 1 | 8 |
| CvixCol | Compactness29 | WD | 3 | 80 | 9,4 | -1 | 7 |
| CvixCol | Compactness29 | WD | 4 | 21 | 2,9 | 1 | 2 |
| CvixCol | Compactness29 | WD | 5 | 86 | 7,8 | -1 | 4 |
| CvixCol | Compactness29 | WD | 5 | 36 | 2,5 | 1 | 1 |
| CvixCol | Compactness29 | WD | 5 | 55 | 2,6 | 1 | 2 |
| CvixCol | Compactness29 | WW | 1 | 29 | 2,5 | 1 | 2 |
| CvixCol | Compactness29 | WW | 2 | 47 | 11,6 | 1 | 8 |
| CvixCol | Compactness29 | WW | 3 | 5 | 3 | -1 | 2 |
| CvixCol | Compactness29 | WW | 3 | 80 | 9,4 | -1 | 7 |
| CvixCol | Compactness29 | WW | 4 | 15 | 3,5 | 1 | 3 |
| CvixCol | Compactness29 | WW | 4 | 32 | 2,8 | 1 | 4 |
| CvixCol | Compactness29 | WW | 5 | 92 | 8,4 | -1 | 4 |
| CvixCol | Compactness29 | WW | 5 | 55 | 2,6 | 1 | 2 |

**Supplementary Table S4:**
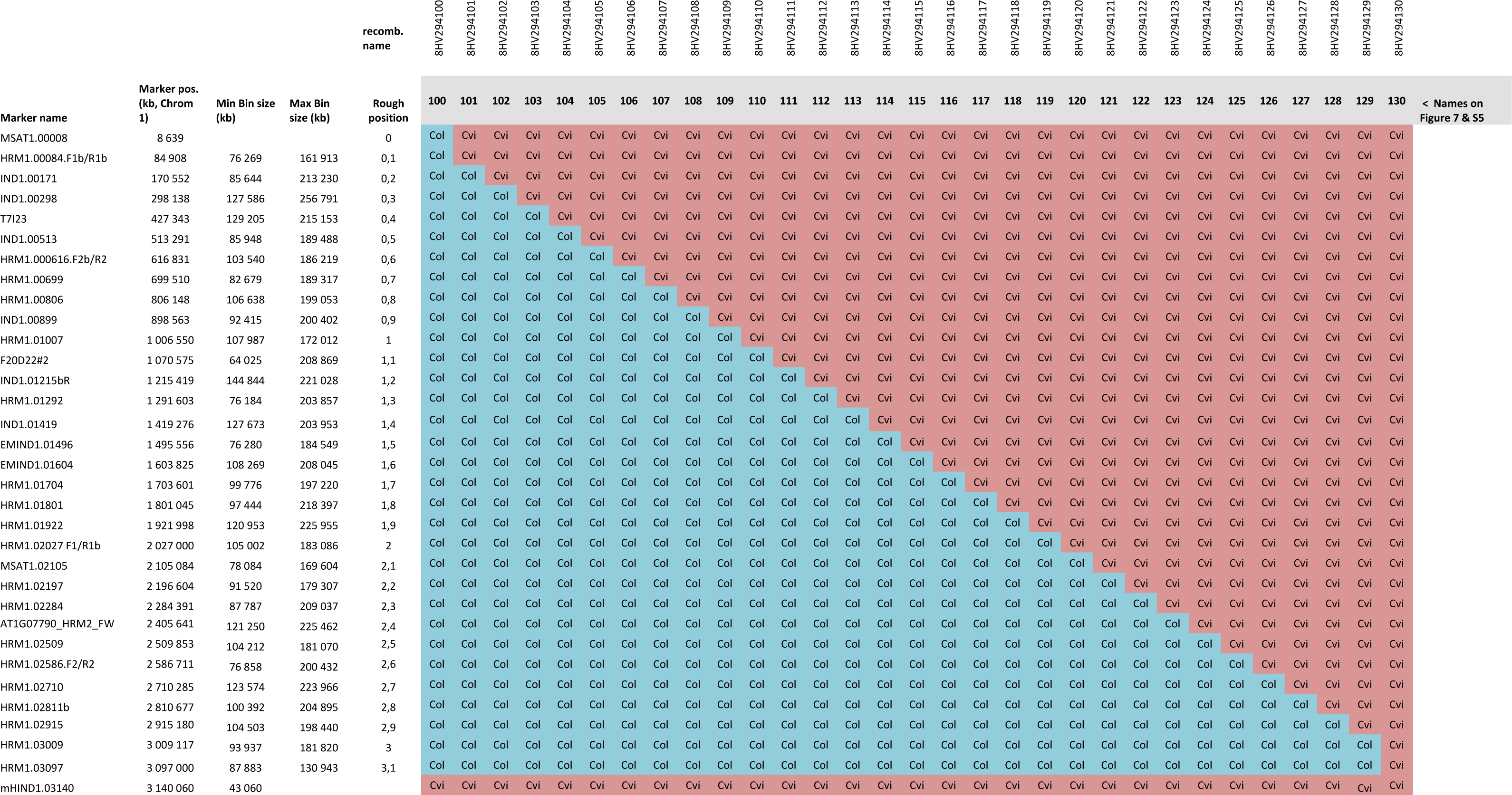
microStairs; genotypic description of recombinant lines. The genotypes of the 31 lines selected for the microStairs experiment are indicated along this 3Mb region of the top of chromosome 1 with markers localised on the physical map (kb).

**Supplementary Table S5:**
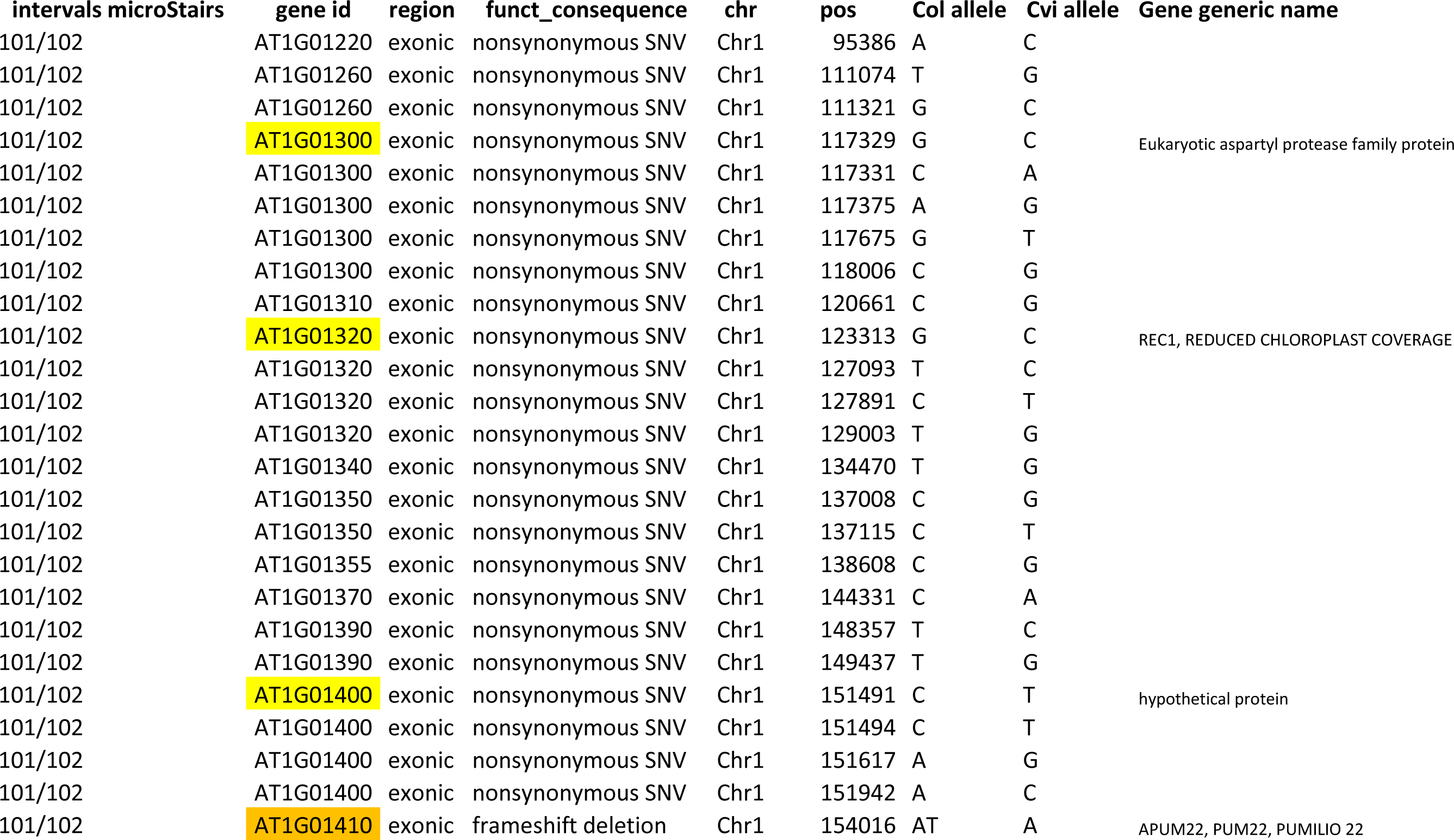
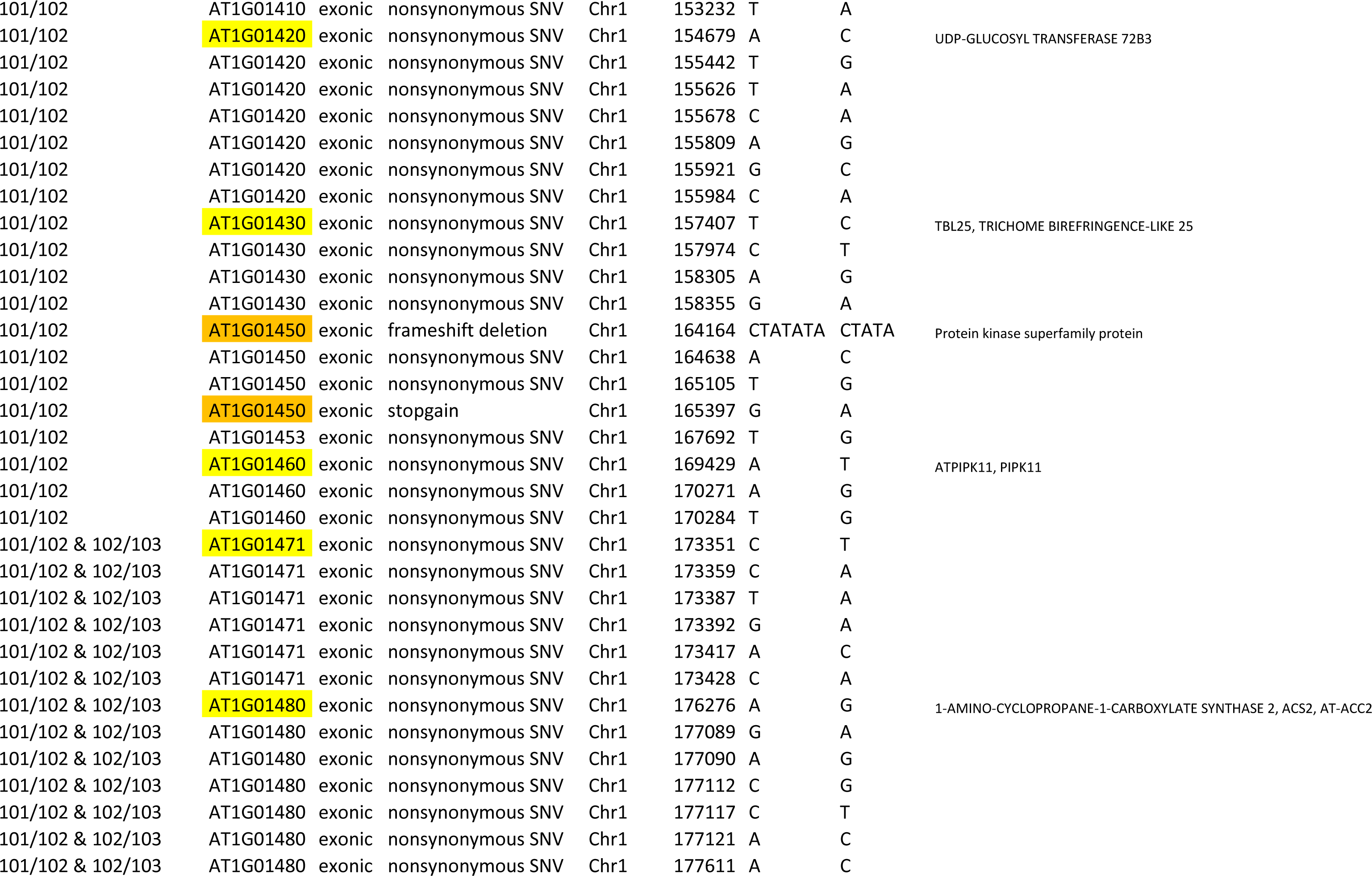
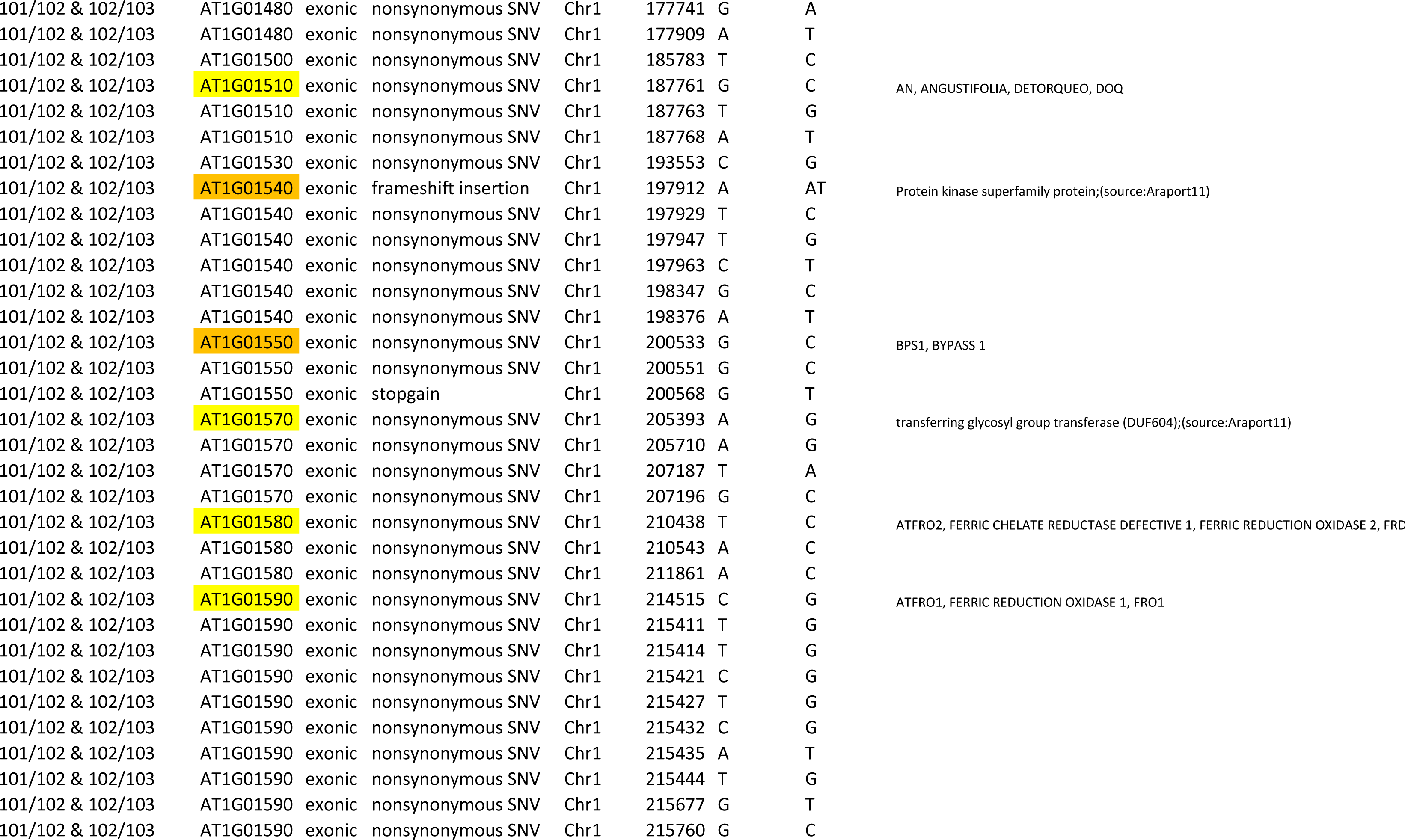
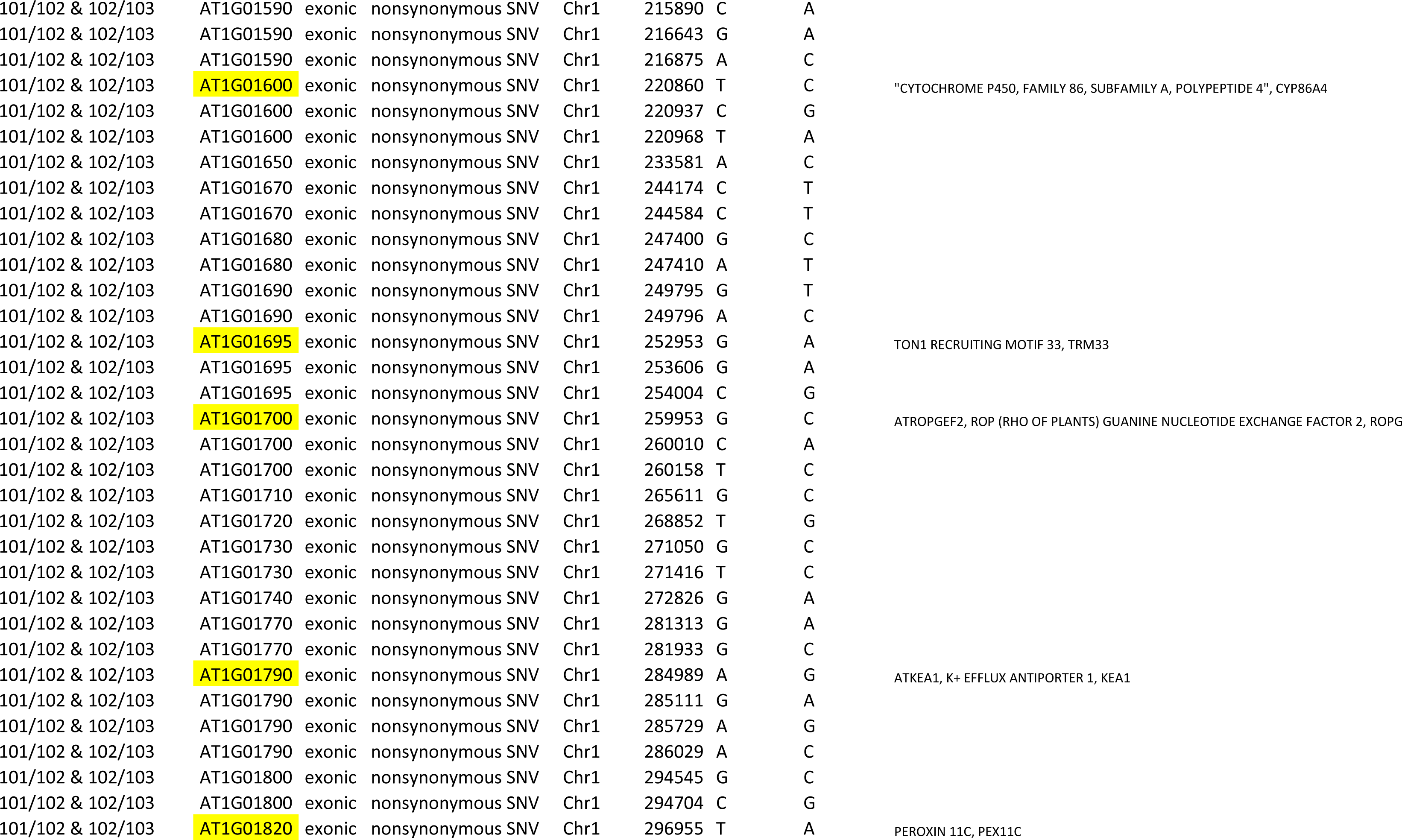
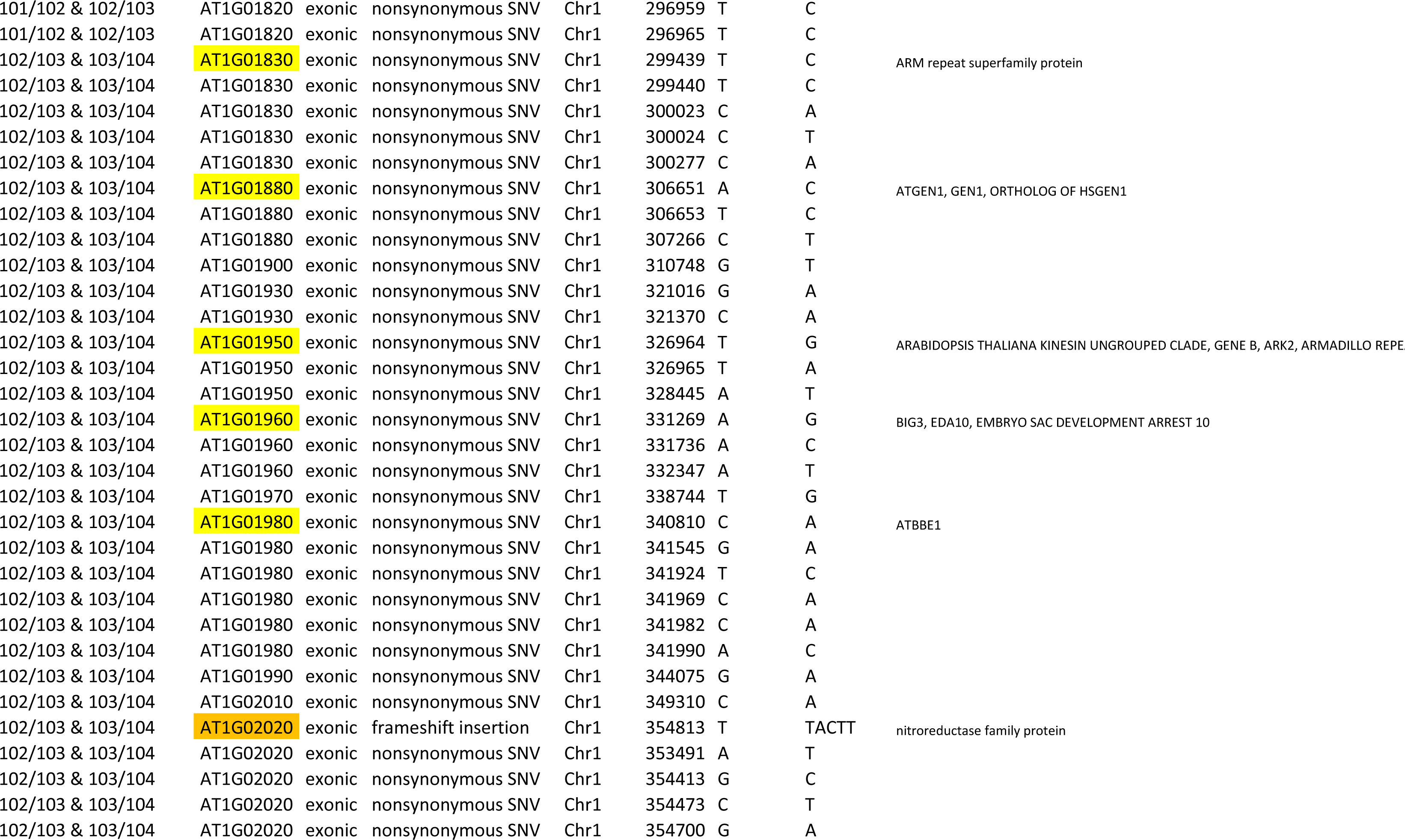
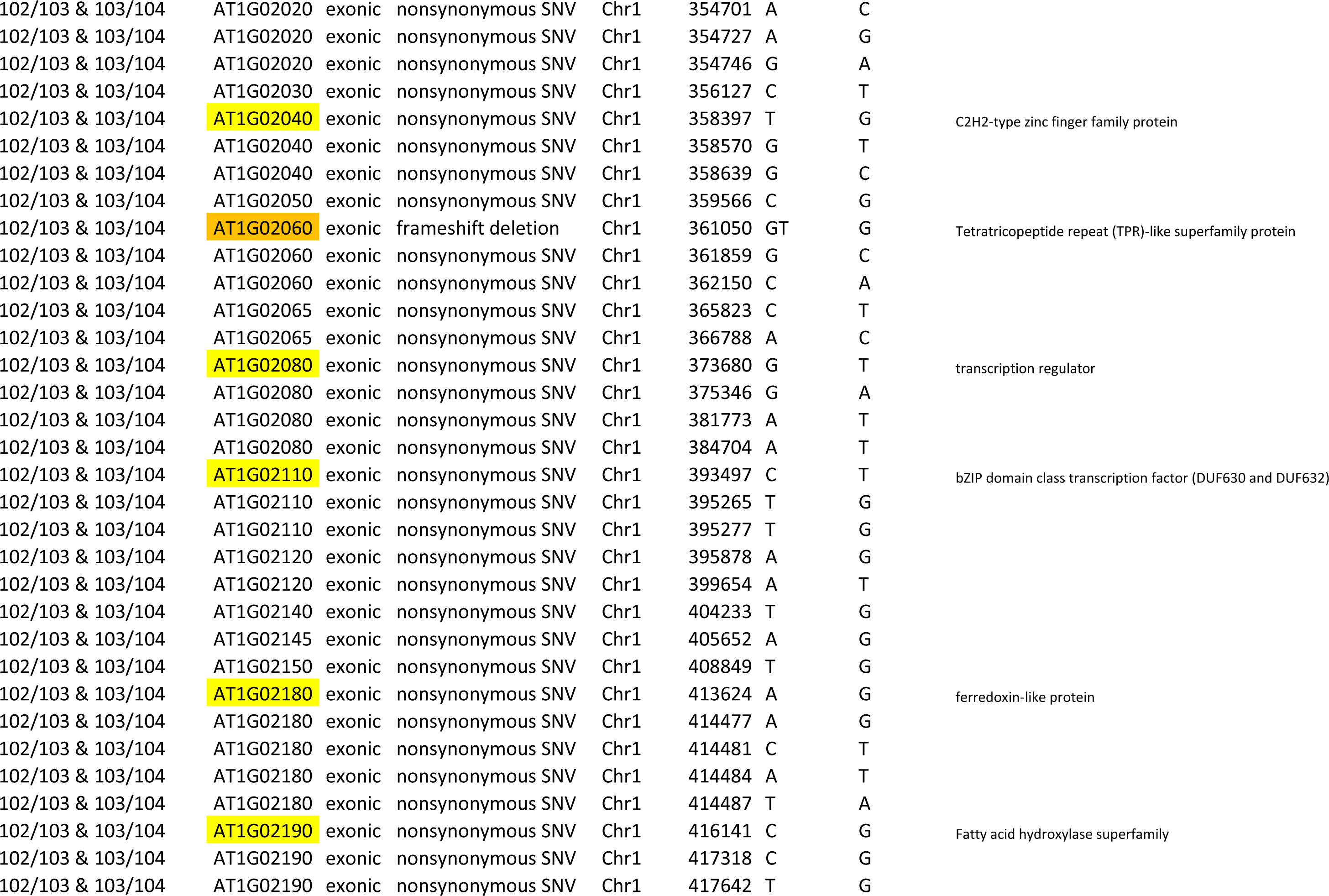
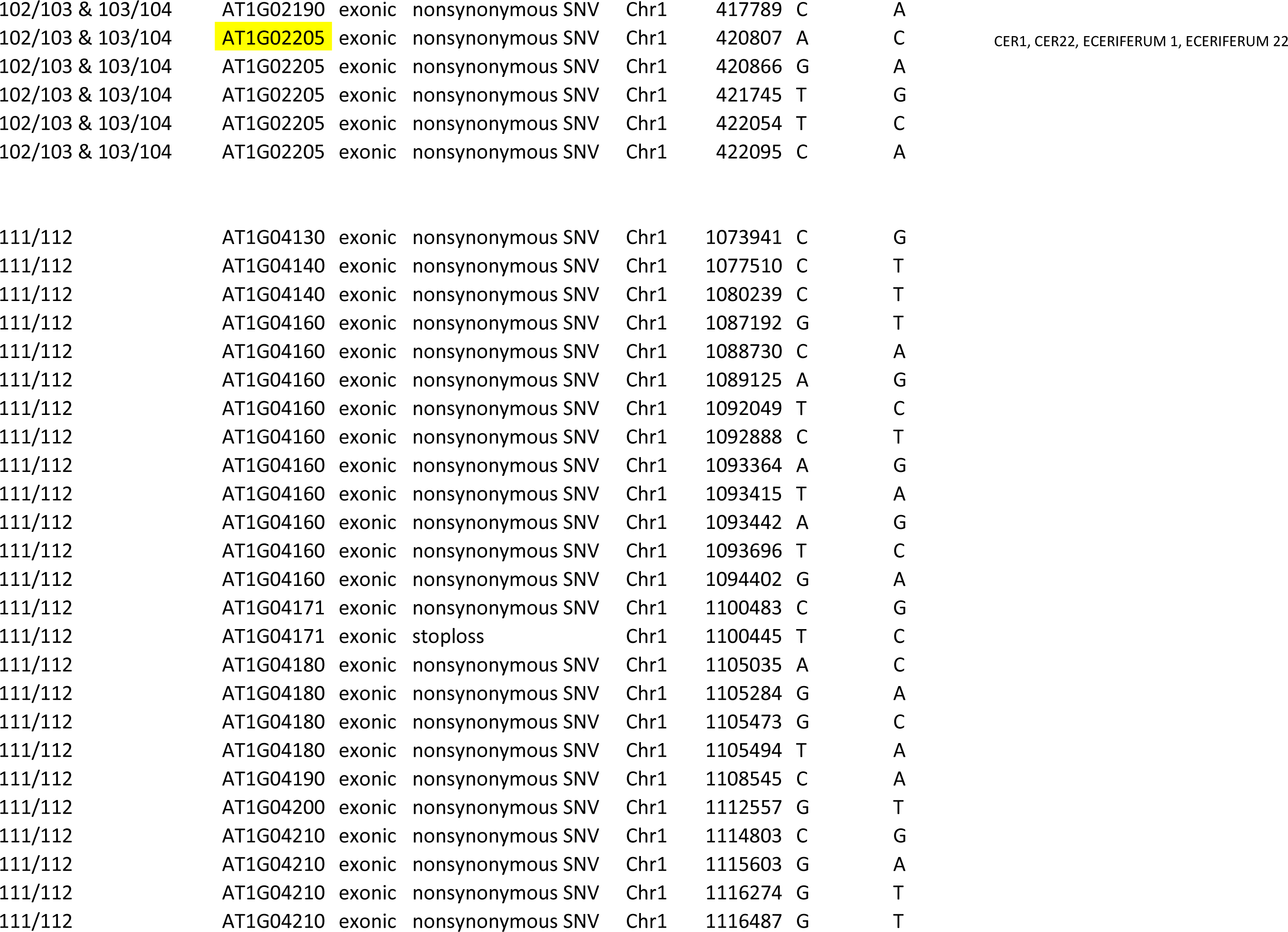
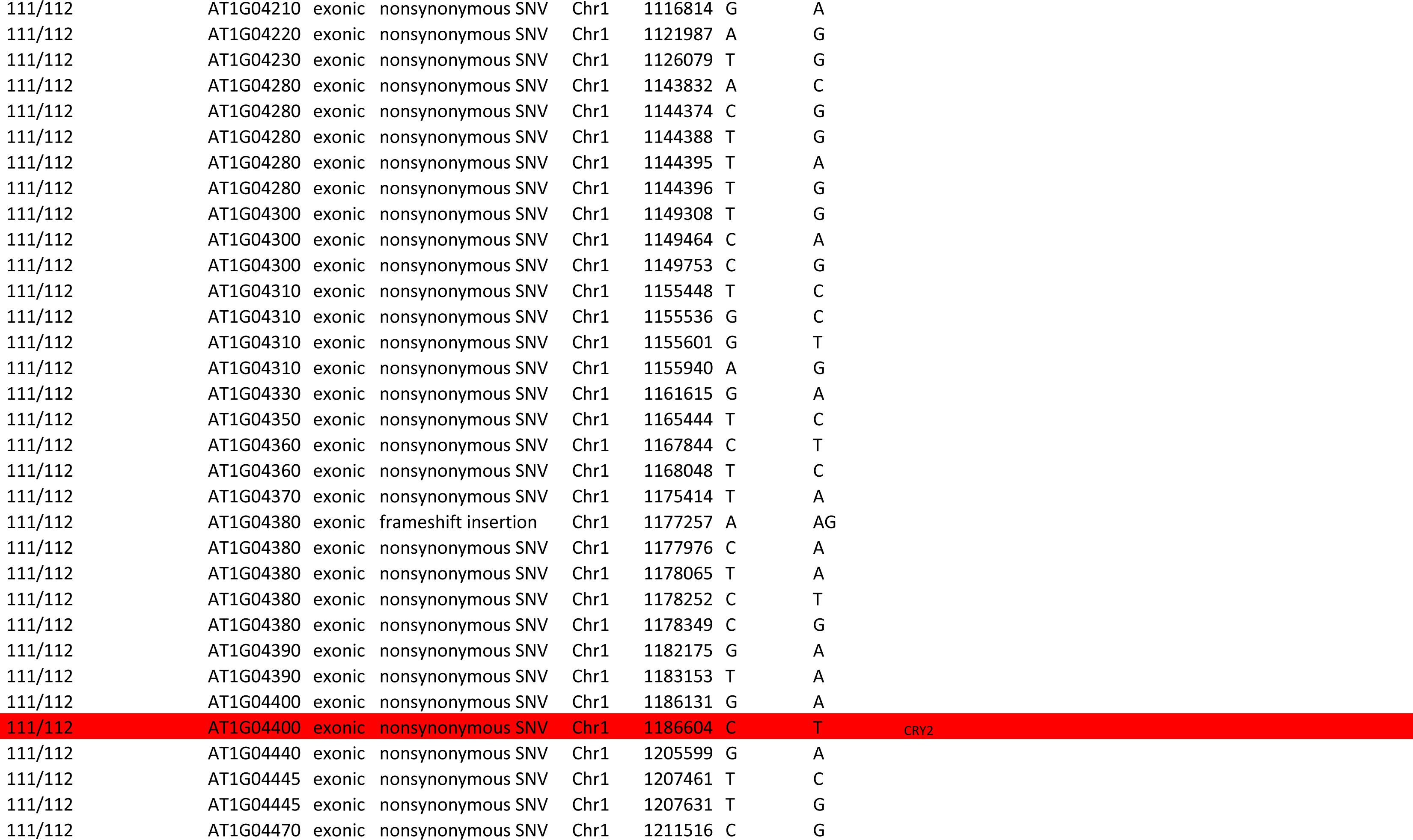
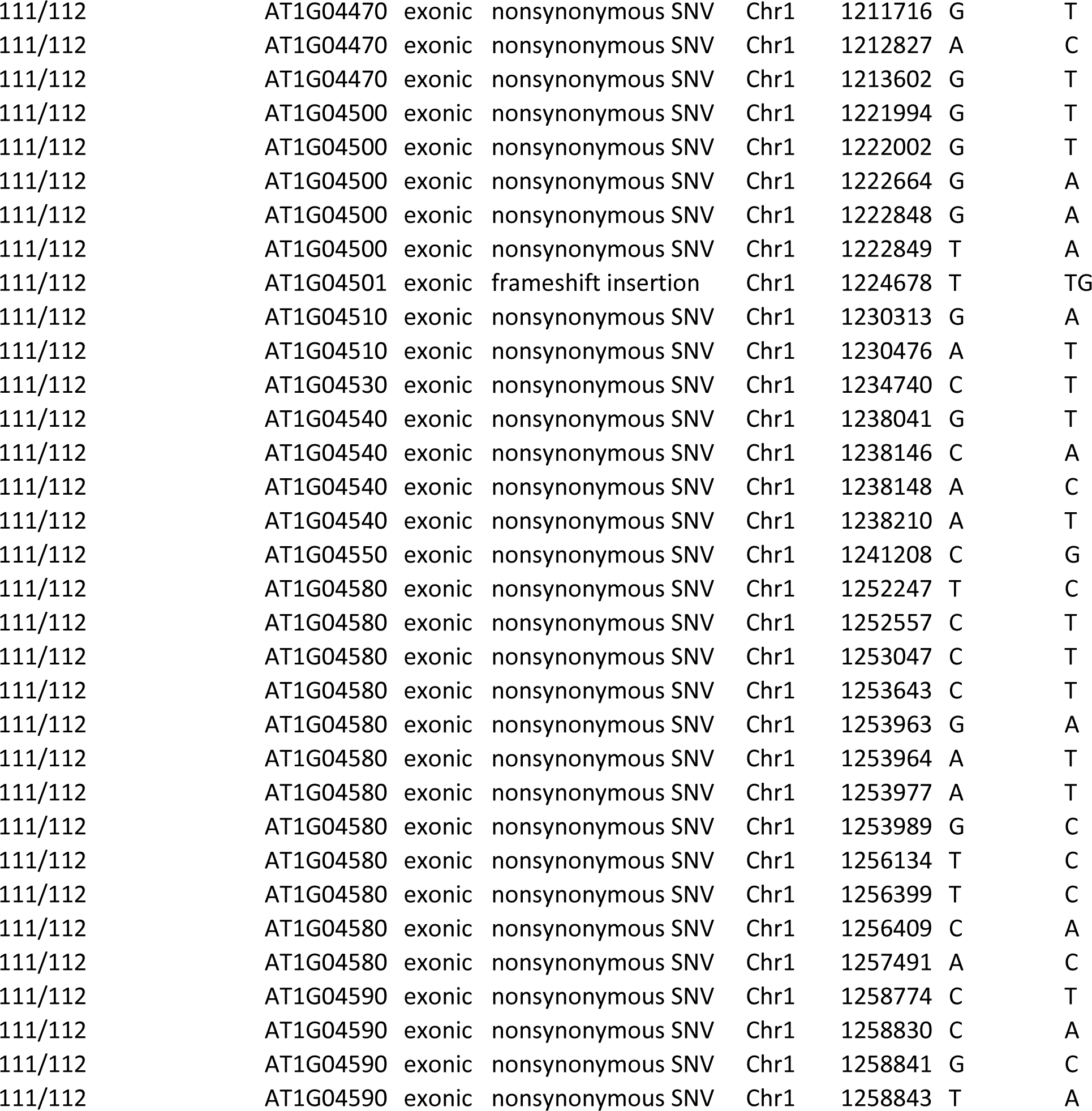
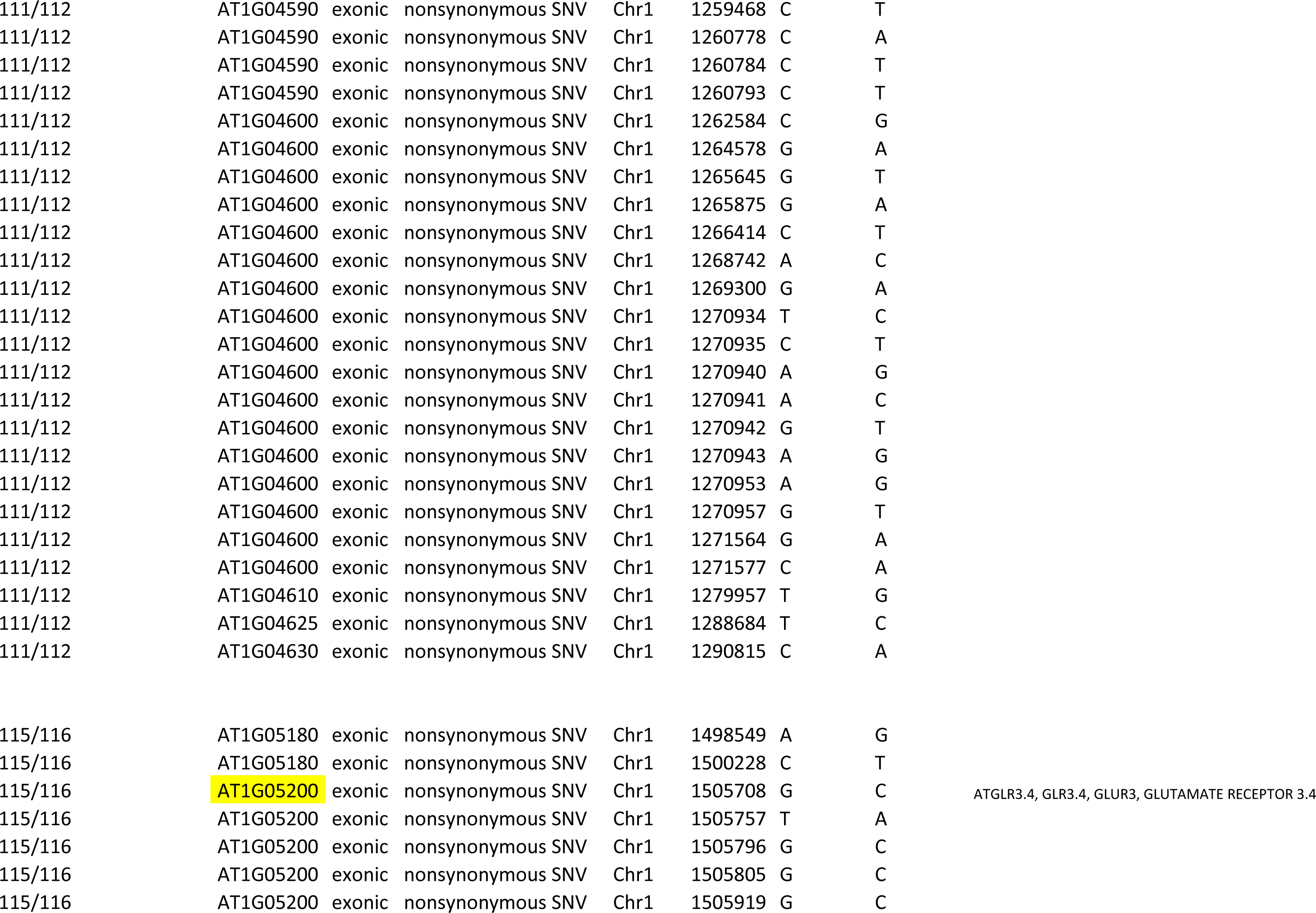
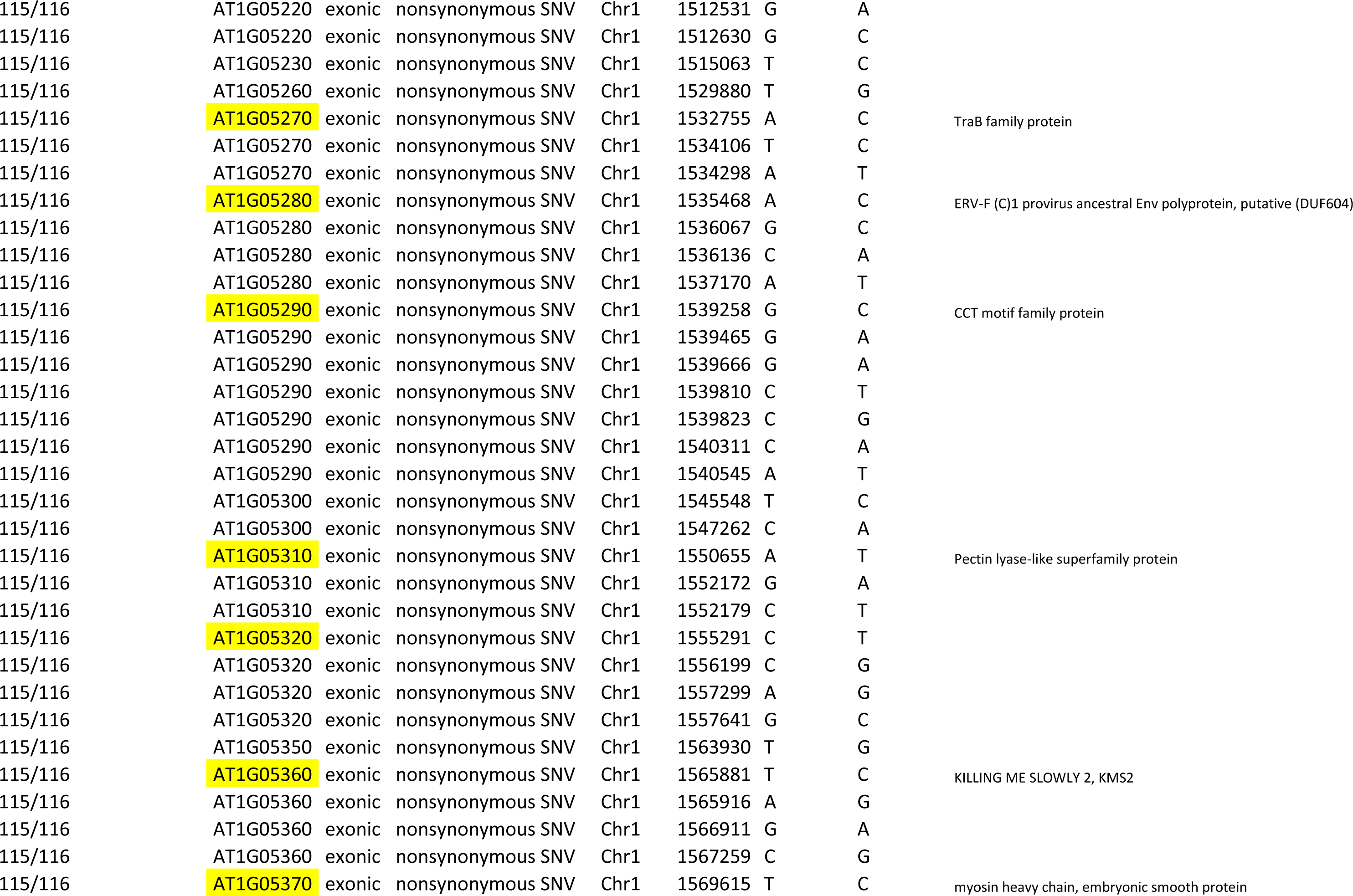
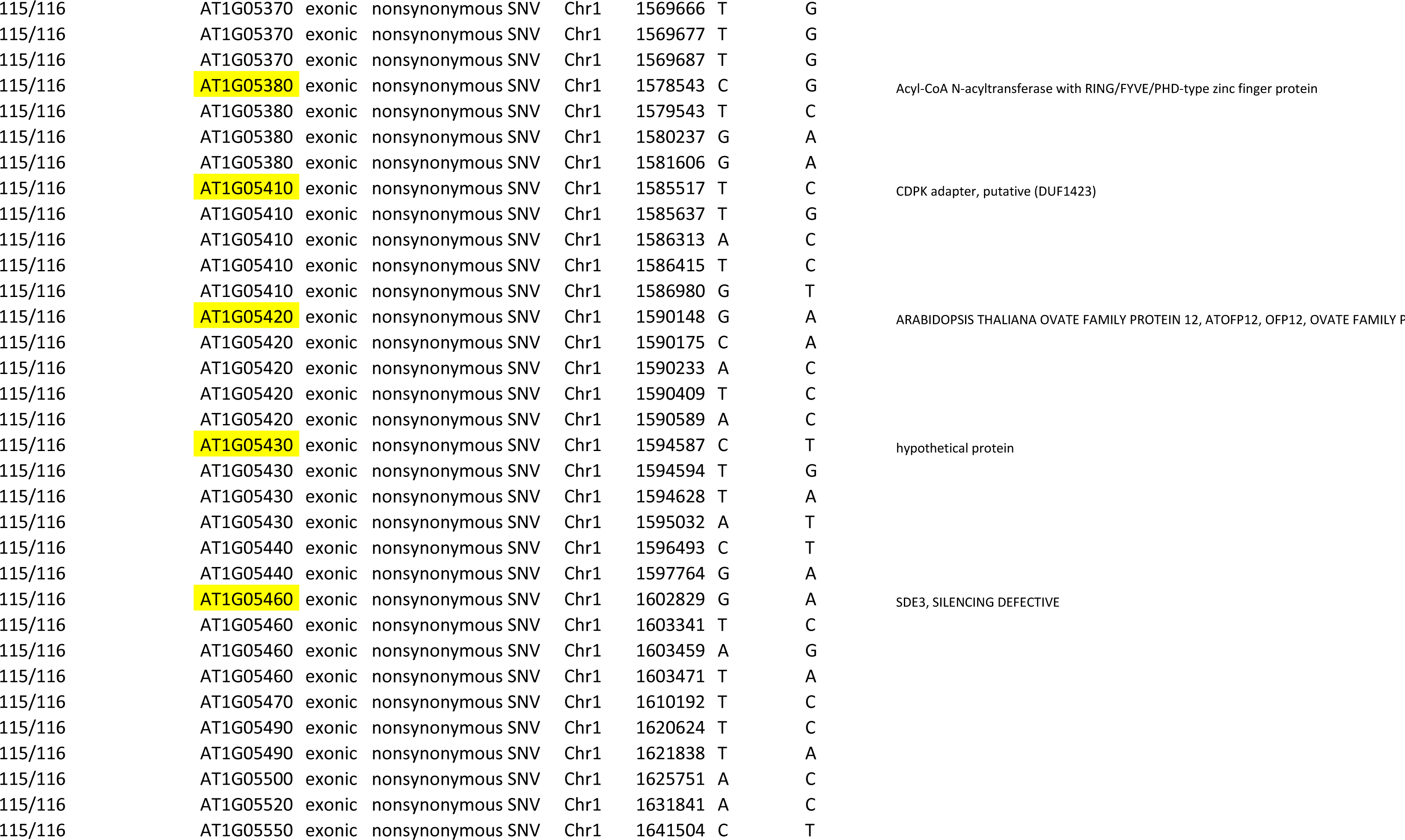
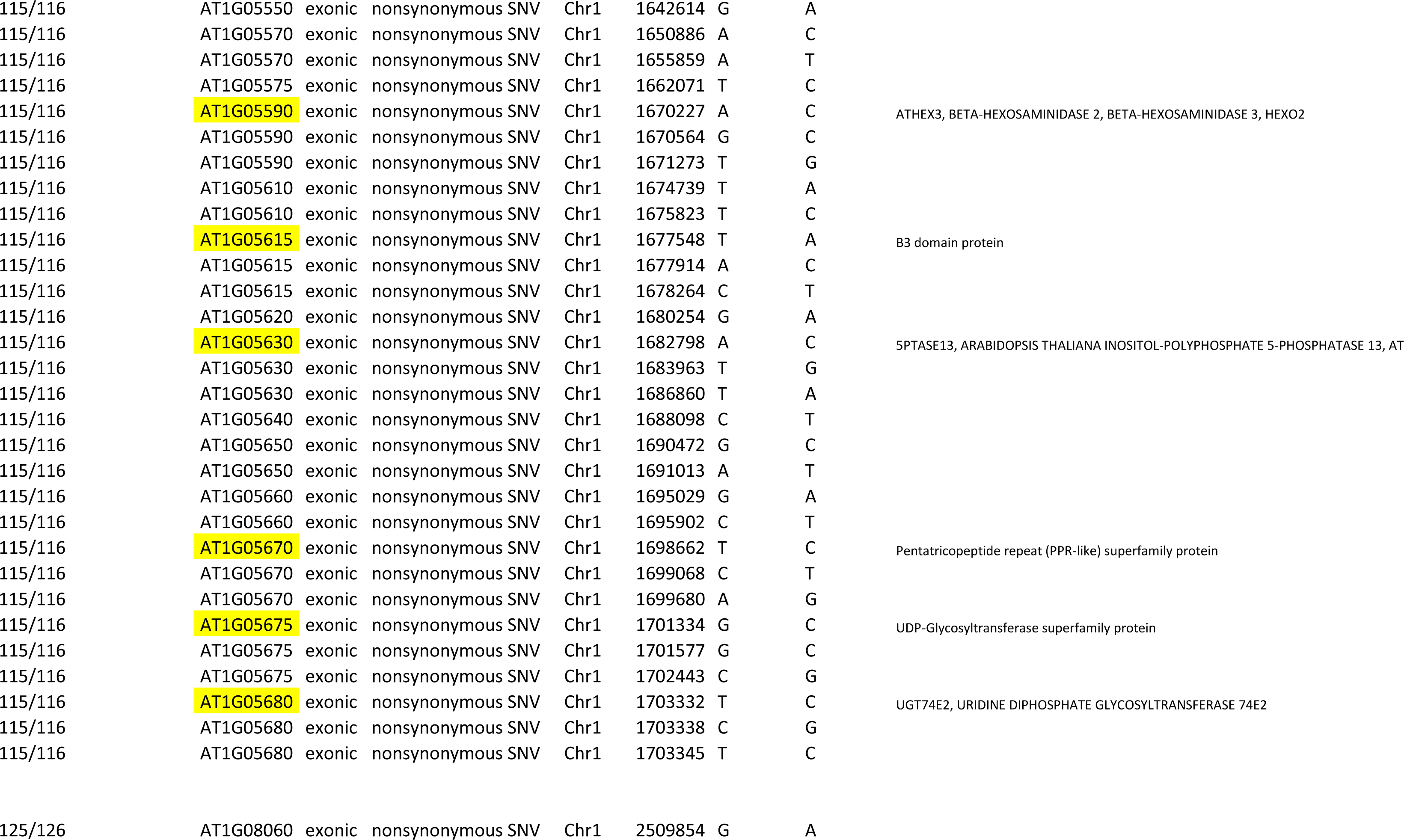
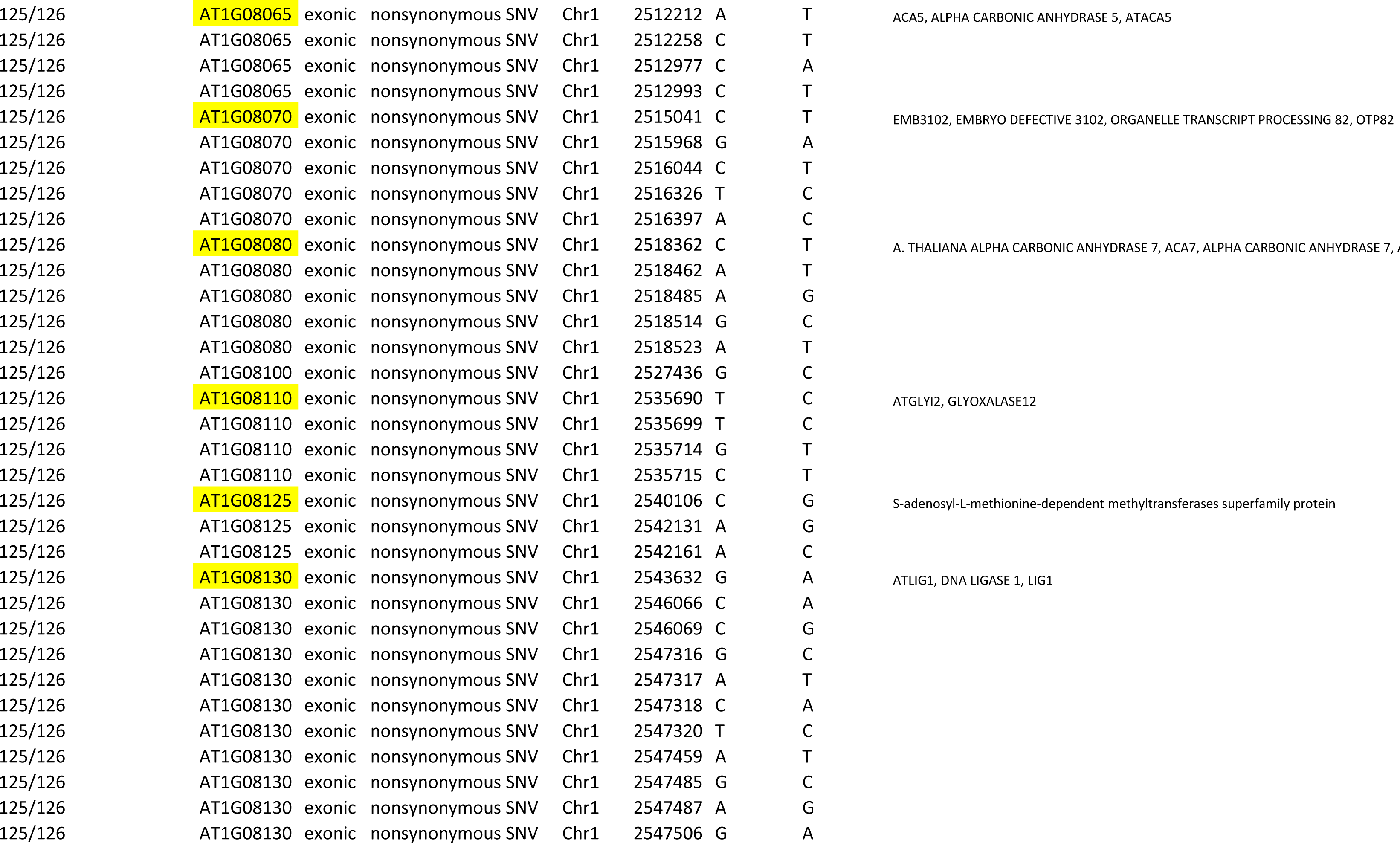
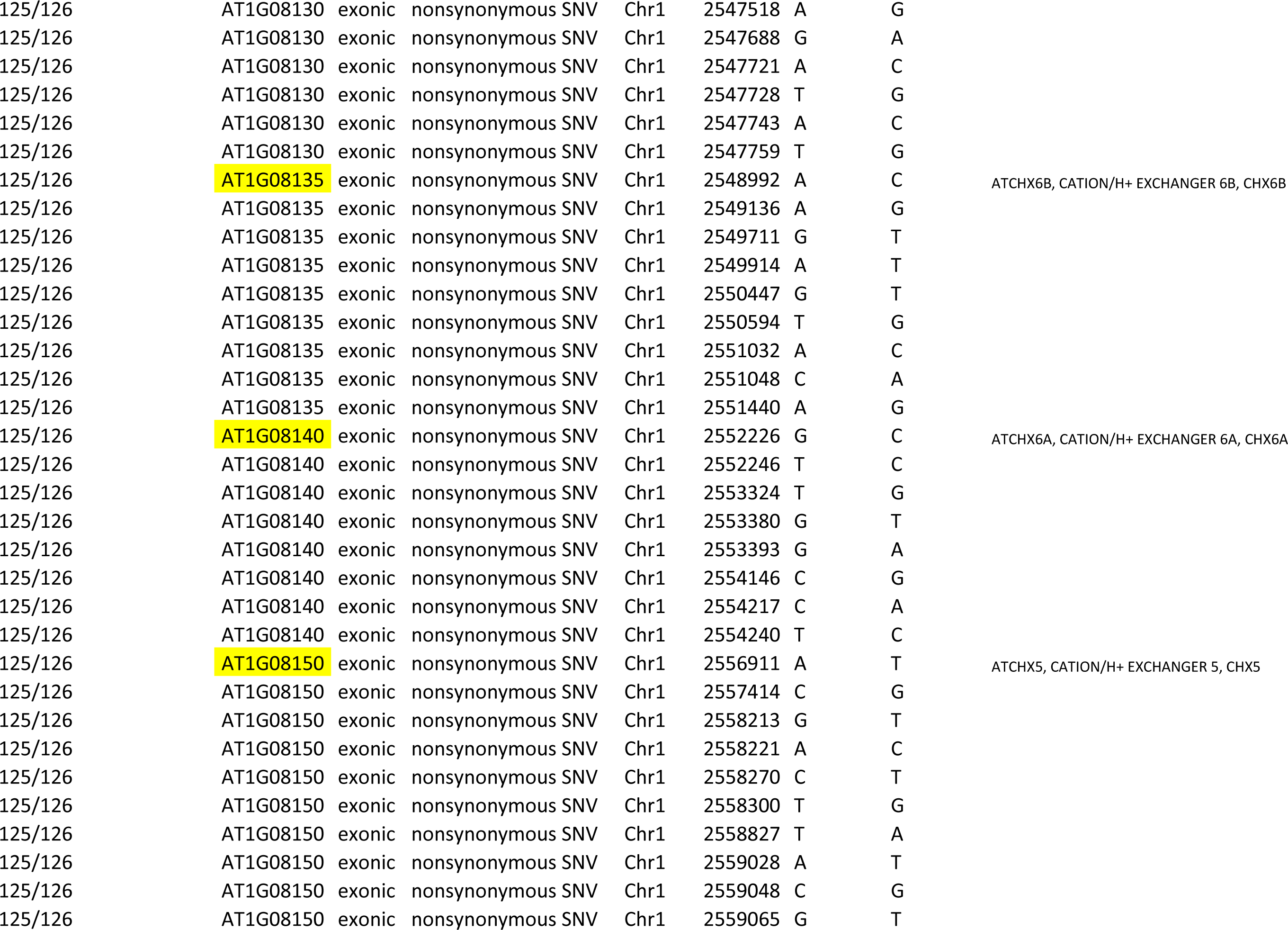
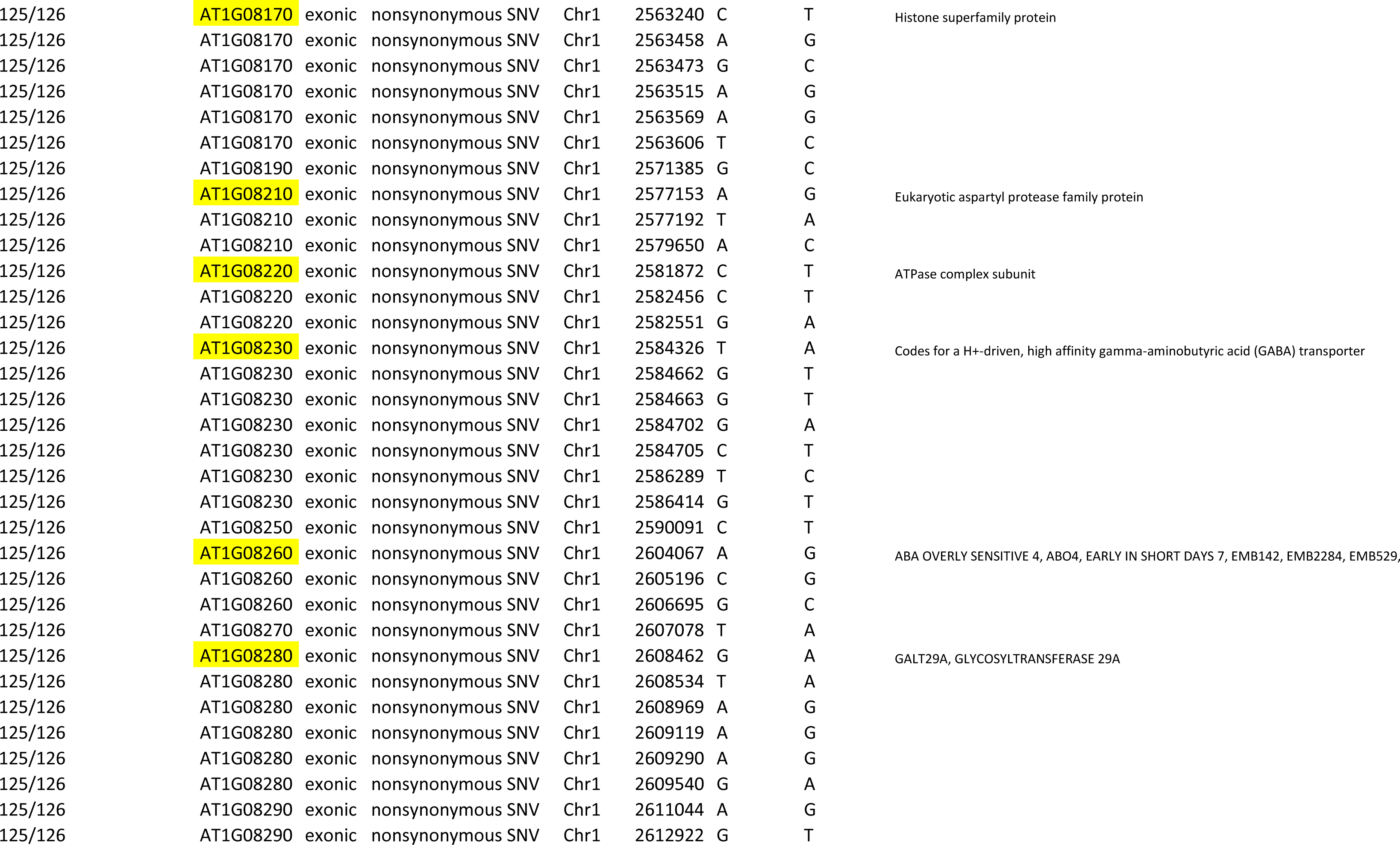
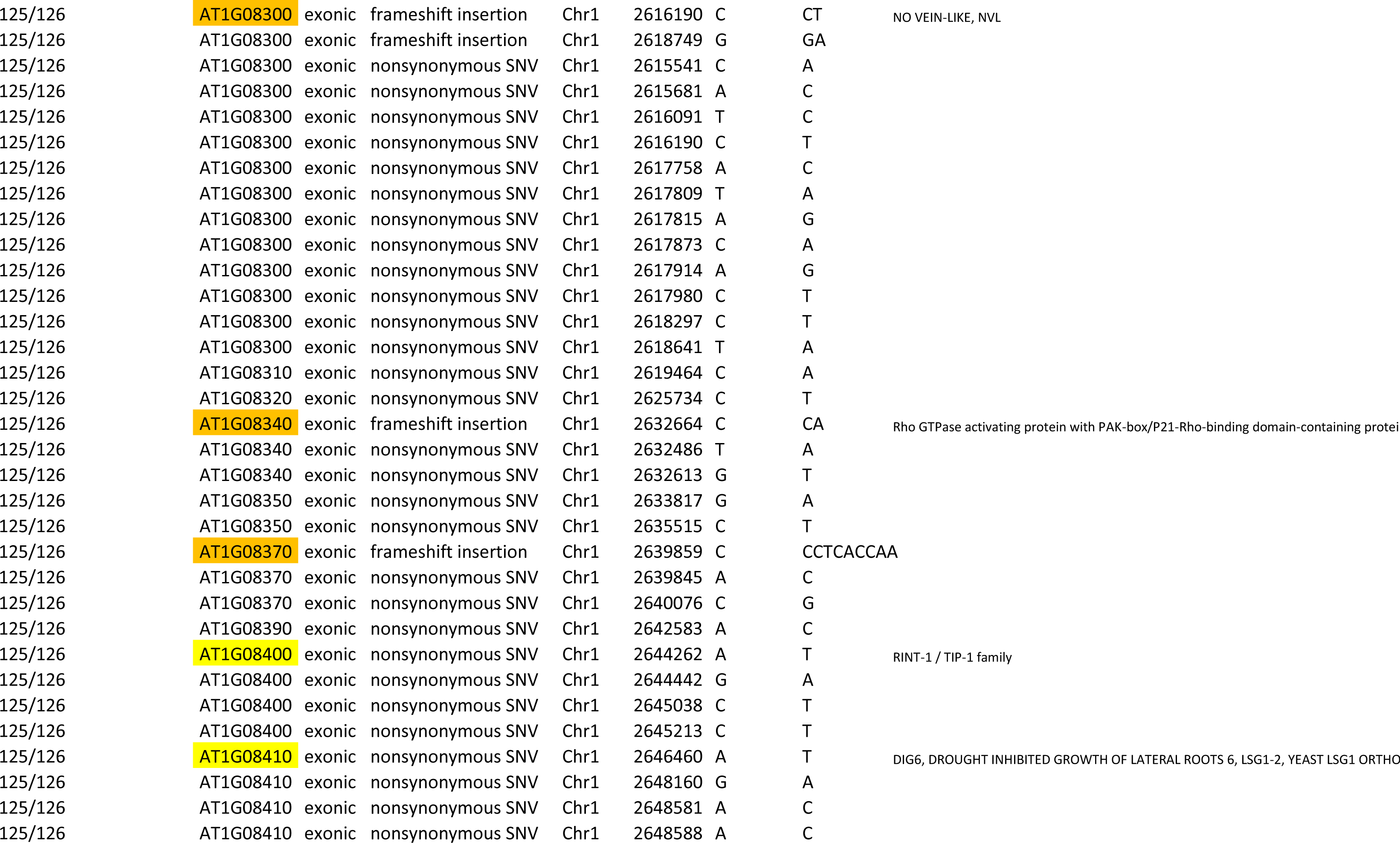
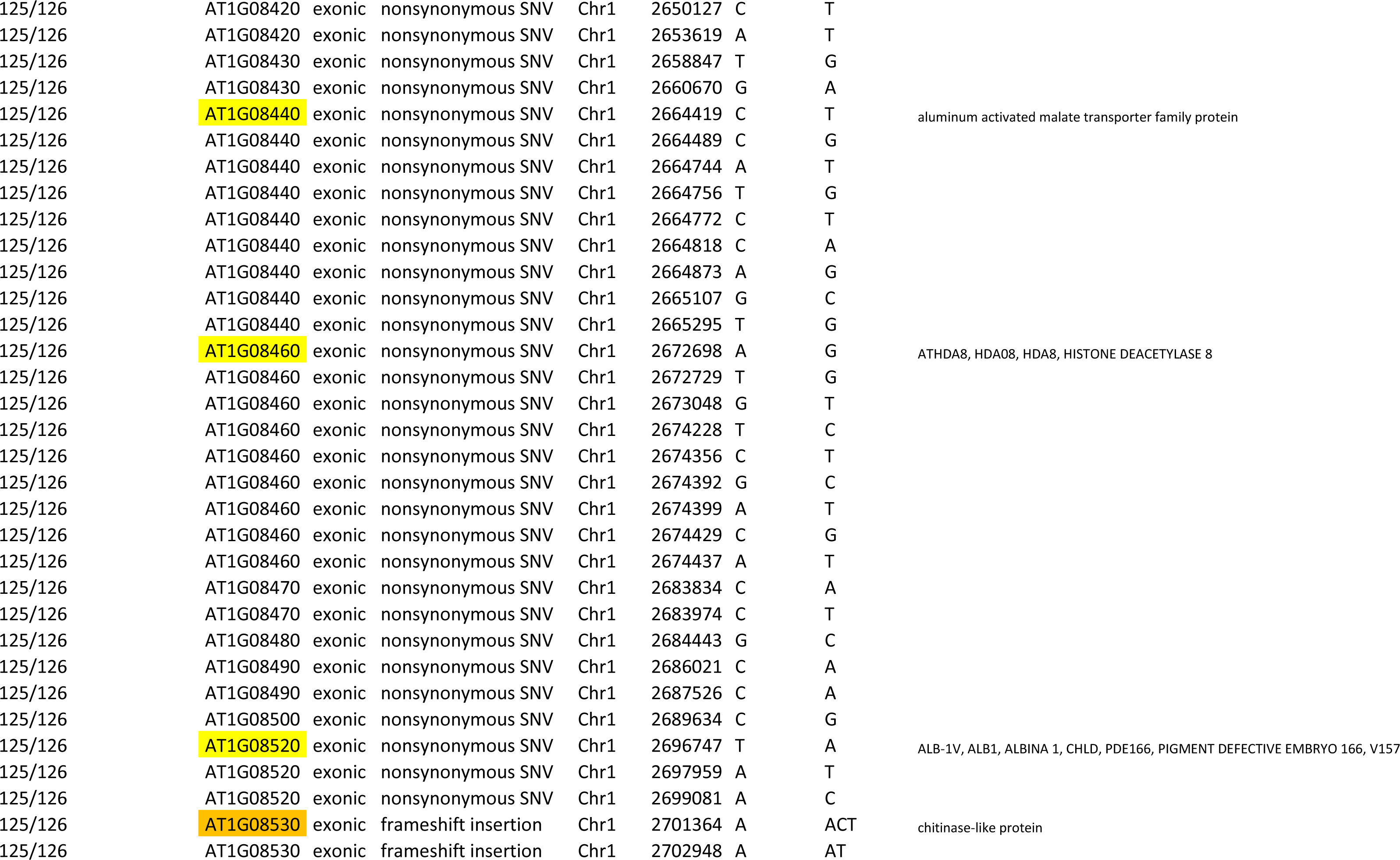
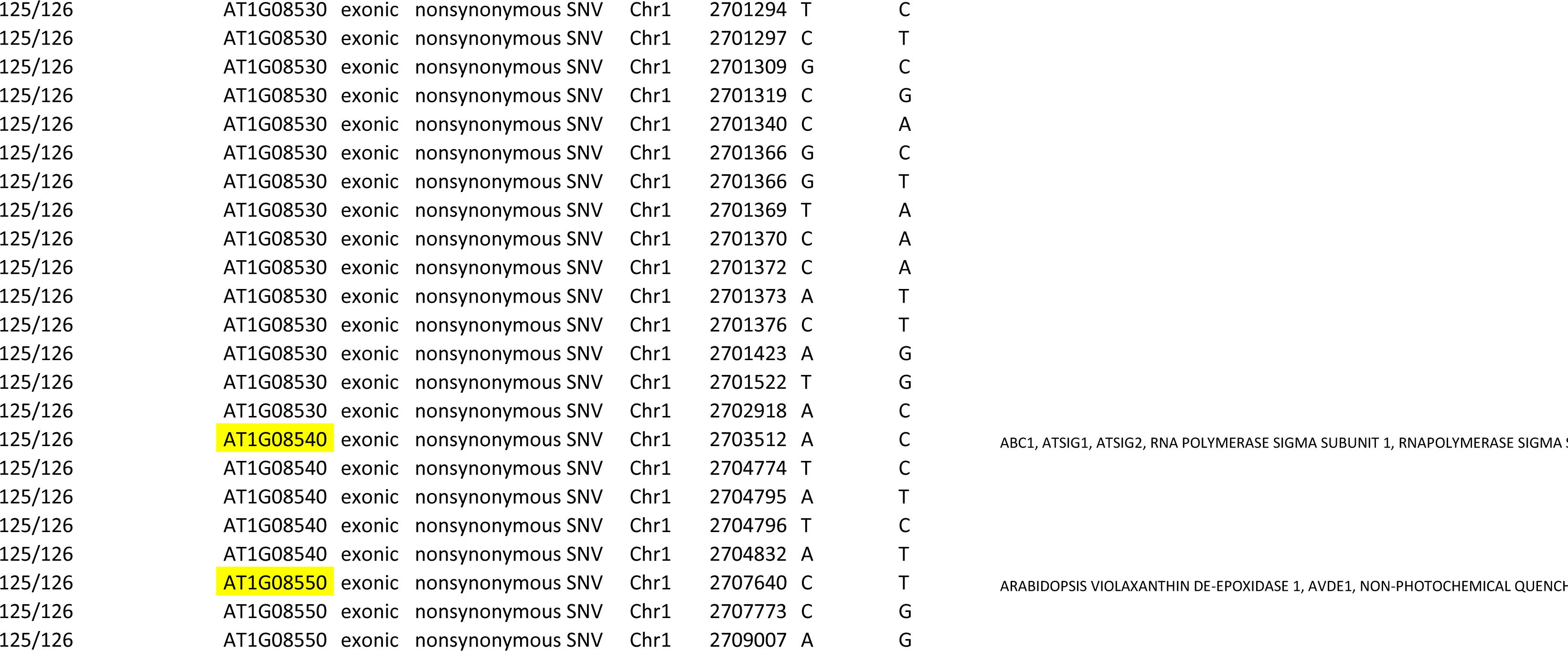
microStairs; polymorphisms segregating in different bins. This table lists all known polymorphisms between Col-0 and Cvi-0 that can potentially affect gene function (according to 1001Genomes data) within PRA29-significant bins only: in the selected physical interval, genes with more than 3 non-synonymous SNP or a high-impact polymorphism are highlighted in yellow and orange respectively. CRY2 candidate gene is highlighted in red.

**Supplementary Table S6:**
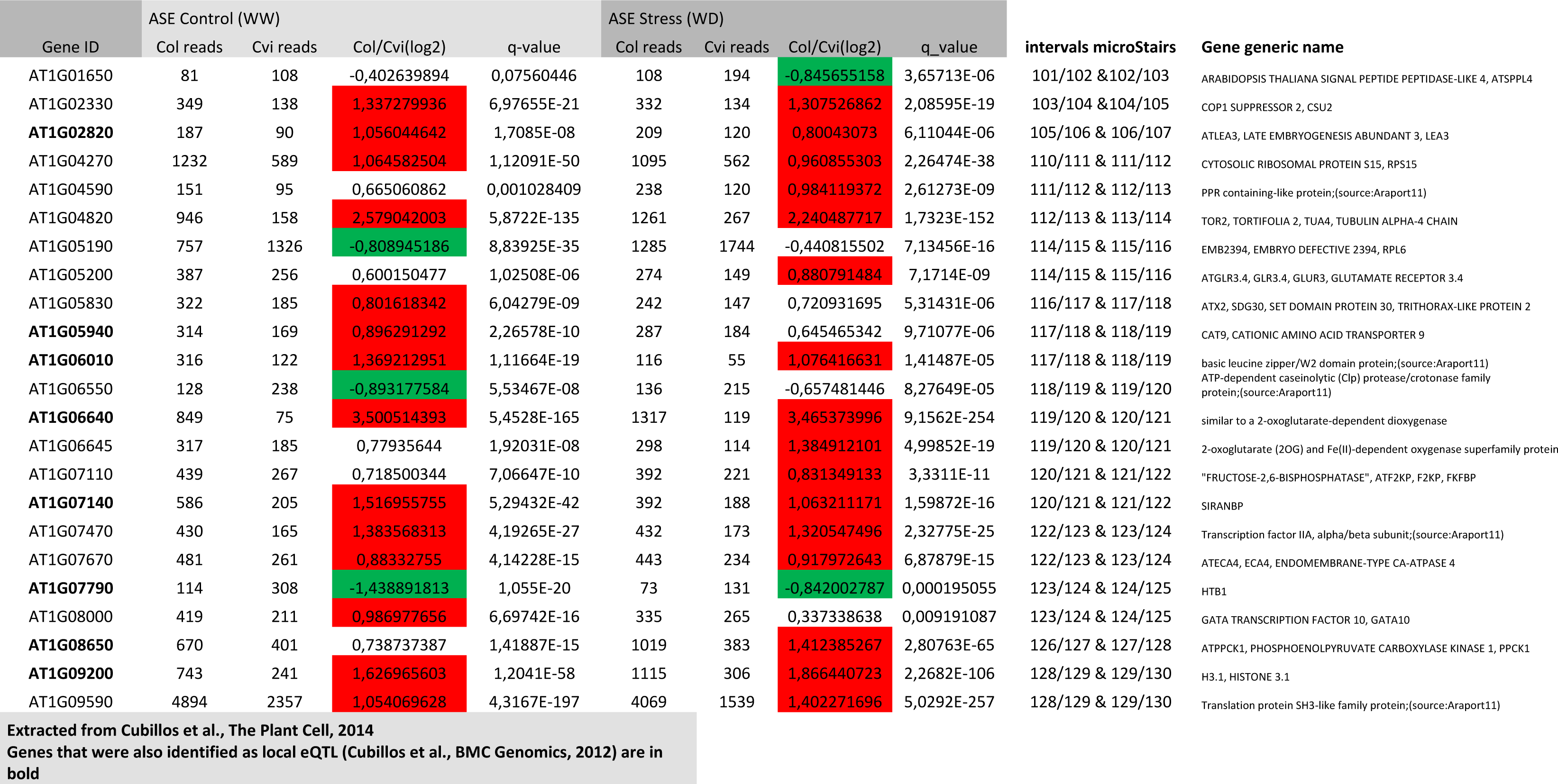
microStairs; differentially expressed genes controlled in cis. List of Cvi/Col differentially expressed genes in an allele-specific fashion ([log2]>0.8 in at least one condition; from Cubillos *et al.*, 2014) across the whole region.

